# *PITAR*, a DNA damage-inducible Cancer/Testis long noncoding RNA, inactivates p53 by binding and stabilizing TRIM28 mRNA

**DOI:** 10.1101/2023.04.11.536370

**Authors:** Samarjit Jana, Mainak Mondal, Sagar Mahale, Bhavana Gupta, Kaval Reddy Prasasvi, Lekha Kandasami, Neha Jha, Abhishek Chowdhury, Vani Santosh, Chandrasekhar Kanduri, Kumaravel Somasundaram

## Abstract

In tumors with WT p53, alternate mechanisms of p53 inactivation are reported. Here, we have identified a long noncoding RNA, PITAR (*p*53 *I*nactivating *T*RIM28 *a*ssociated *R*NA), as an inhibitor of p53. PITAR is an oncogenic Cancer/testis lncRNA and is highly expressed in glioblastoma (GBM) and glioma stem-like cells (GSC). We establish that TRIM28 mRNA, which encodes a p53-specific E3 ubiquitin ligase, is a direct target of PITAR. PITAR interaction with TRIM28 RNA stabilized TRIM28 mRNA, which resulted in increased TRIM28 protein levels and reduced p53 steady-state levels due to enhanced p53 ubiquitination. DNA damage activated PITAR, in addition to p53, in a p53-independent manner, thus creating an incoherent feedforward loop to inhibit the DNA damage response by p53. While PITAR silencing inhibited the growth of WT p53 containing GSCs in vitro and reduced glioma tumor growth in vivo, its overexpression enhanced the tumor growth in a TRIM28-dependent manner and promoted resistance to Temozolomide. Thus, we establish an alternate way of p53 inactivation by PITAR, which maintains low p53 levels in normal cells and attenuates the DNA damage response by p53. Finally, we propose PITAR as a potential GBM therapeutic target.

## Introduction

Much of the human genome once considered as “junk DNA” is now shown to be pervasively transcribed into thousands of long noncoding RNAs (lncRNAs). Accumulated evidence over the last two decades implicated lncRNA in fundamental biological processes that play an essential role in several pathological conditions like cancer (Djebali *et al*., 2012; Slack and Chinnaiyan, 2019a). LncRNA constitutes a major subset among ncRNAs and is arbitrarily defined as transcripts of longer than 500 bps, which are spliced, 5’ capped, and poorly conserved across species (Cabili *et al*., 2011; Derrien *et al*., 2012; Iyer *et al*., 2015; Mattick *et al*., 2023). LncRNAs function through multiple mechanisms, such as their specific interactions with DNA, RNA, and proteins, that can modulate chromatin structure, regulate the assembly and function of membrane-less bodies, alter the stability and translation of mRNAs and interfere with their signaling pathways (Statello *et al*., 2020).

The tumor suppressor protein p53 plays an important role in preserving genome integrity and inhibiting malignant transformation (Levine, 1997). p53, which gets activated in response to stress like DNA damage, activates the transcription of many protein coding genes that control several cellular processes like cell cycle, programmed cell death, and senescence (Riley *et al*., 2008; Beckerman and Prives, 2010; Bieging and Attardi, 2012). p53 is mutated in more than 50% of human cancers (Vogelstein, Lane and Levine, 2000; Vousden and Lane, 2007). In cancers that do not carry mutations in p53, the inactivation occurs through other genetic or epigenetic alterations (Olivier *et al*., 2002; Vousden and Lu, 2002; Mitra *et al*., 2021).

LncRNAs have been shown to play an important role in the p53 network. While several lncRNAs such as LincRNA-p21, DINO, PANDA, LINC-PANT, GUARDIN, NEAT1, NBAT1 (Mitra *et al*., 2021), and PVT1 are shown to function as downstream effectors of p53, other lncRNA like MEG3, MALAT1, H19, Linc-ROR, and PSTAR act as upstream regulators of p53 (Jain, 2020). Our study identified a lncRNA called *p*53 *I*nactivating *T*RIM28 *a*ssociated *R*NA (PITAR) as a p53 inactivator with a protumorigenic function. PITAR is highly expressed in glioblastoma (GBM) and glioma stem-like cells (GSCs) and interacts with TRIM28 mRNA, which encodes a p53-specific E3 ubiquitin ligase. TRIM28 inhibits p53 through HDAC1-mediated deacetylation and direct ubiquitination in an MDM2-dependent manner (Wang *et al*., 2005a). PITAR-TRIM28 interaction stabilized TRIM28 mRNA resulting in higher levels of TRIM28 protein that promoted ubiquitin-mediated degradation of p53. We also found that PITAR is essential for glioma tumor growth, and PITAR is induced by DNA damage in a p53-independent manner. Thus, our study discovered PITAR as an inhibitor of p53 via a unique mechanism of interaction with TRIM28 mRNA and a potential target for developing novel therapy.

## Results

### Identification of FAM95B1/PITAR, a conserved cancer/testis lncRNA that promotes cell proliferation in GBM

In a recent study involving our group’s transcriptional profiling of mRNAs and lncRNAs, we identified several GBM-specific clinically relevant lncRNA regulatory networks (Paul *et al*., 2018). To identify GBM-associated lncRNAs with functional relevance to glioma stem-like cells (GSCs) biology, we integrated the differentially regulated RNAs (DEGs) from GBM vs control brain samples (TCGA) (**Supplementary Table 1)** with DEGs from GSC vs differentiated glioma cells (DGCs) (Suvà *et al*., 2014) (**Figure 1A**). The GBM and GSC integrated analysis revealed three interesting lncRNA PVT1, H19, and FAM95B1, showing significant upregulation in both GBM and GSCs (**Figure 1A).** Of the three GBM and GSC-specific regulated lncRNAs, PVT1 and H19 have been extensively investigated for their role in cancer development and progression (Xue *et al*., 2018; Azab and Azzam, 2021; Li *et al*., 2022; Lv *et al*., 2022; Wang *et al*., 2022), whereas FAM95B1 (ENSG00000223839.7, NONHSAG052279.2, Lnc-ANKRD20A3-51) has not been explored for its role in cancer. We have named the FAM95B1 lncRNA as PITAR (***p***53 ***i***nactivating ***T***RIM28-***a***ssociated lnc***R***NA) as we found it has a strong functional association with the well-known tumor suppressor gene p53.

**Figure 1:**
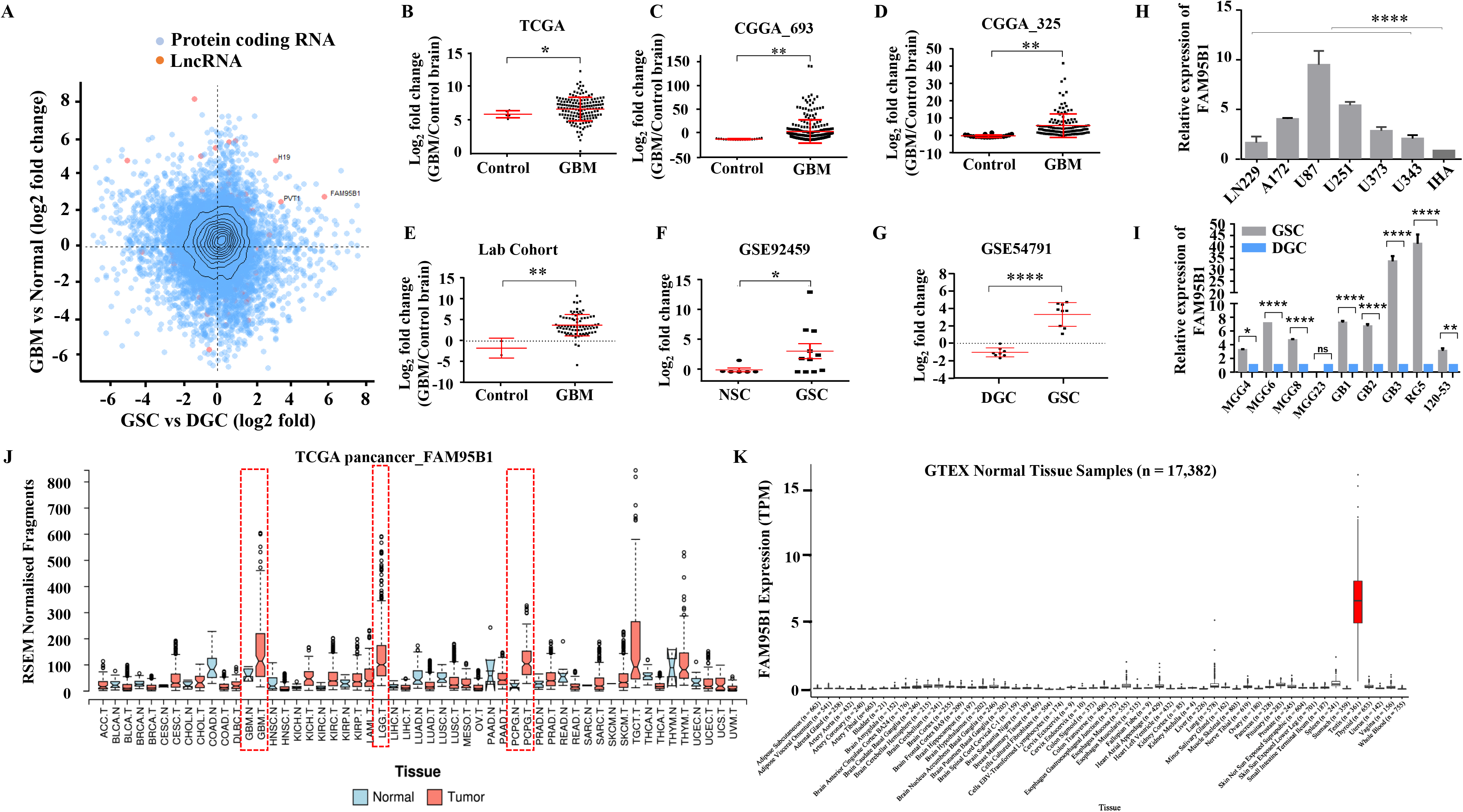
Identifying Glioblastoma stem cell-specific and conserved cancer/testis LncRNA. **A.** The Scatterplot depicts the differentially regulated protein-coding (Blue dots) and lncRNA (Red dots) transcripts. The X-axis shows a differentially regulated gene (log2 fold change) in GBM vs. normal (TCGA patient cohort), and the Y-axis represents differentially regulated genes (log2 fold change) in the GSC Vs. DGC dataset (GSE54791). **B**, **C & D.** The expression in log2 fold change of FAM95B1 was shown in TCGA and two patient cohorts of the CGGA dataset (CGGA_693 & CGGA_325). GlioVis was used to obtain the gene expression matrix, and a t-test was performed using GraphPad Prism v6. **E.** Plot depicts log2 fold change of FAM95B1 in our patient cohort (normal, n=3 and GBM, n=79). **F.** Expression of FAM95B1 in GSC vs. NSC dataset (GSE92459). **G.** Expression of FAM95B1 in GSC vs. DGC dataset (GSE54791). **H.** Relative expression of FAM95B1 was quantified by qRT-PCR in different Glioblastoma cell lines and immortalized human astrocytes (IHA). **I.** Relative expression of FAM95B1 was measured in seven GSCs and its corresponding DGCs using the qRT-PCR method. **J.** Relative expression (RPKM) of FAM95B1 across different cancer types in the TCGA Pan-cancer cohort. **K.** Expression in TPM of FAM95B1 amongst the GTEx normal bulk tissue RNA-seq dataset. Data are shown as mean ± SD (n=3). \*\*\**P*-value <0.001, \*\**P*-value <0.01, \**P*-value <0.05.

PITAR is significantly upregulated in multiple GBM cohorts (**Figure 1B-E**) and patient-derived GSCs, (**Figure 1F and G**). The PITAR transcript also showed a one-to-one significant negative correlation between GSCs, and DGCs (**Supplementary figure 1A).** PITAR promoter harbored active histone marks (H3k27Ac) in GSCs but not in DGCs (**Supplementary figure 1B**); thus, H3K27ac levels at the PITAR promoter in GSCs correlate with its expression status. Additional independent validation confirmed its higher expression in multiple glioma cell lines compared to immortalized astrocytes (**Figure 1H**) and in multiple patient-derived GSC lines (**Figure 1I)**. Pan cancer data analysis revealed specific upregulation of PITAR in GBM and low-grade glioma (LGG) except in neuroendocrine tumor Pheochromocytoma and Paraganglioma (PCPG) (**Figure 1J**). Further, the interrogation of the Genotype-Tissue Expression (GTEx) of normal tissues revealed a highly tissue-restricted expression of PITAR in the testis, thus resembling the expression properties of a cancer-testis antigen (**Figure 1K**).

PITAR, located in human chromosome 9, has a 4236-nucleotide-long transcript (Transcript: ENST00000455995.5) with four exons (**Supplementary** Figure 2A). The noncoding nature of PITAR was confirmed by the Coding Potential Assessing Tool (CPAT) (https://www.ncbi.nlm.nih.gov/orffinder/; data not shown). The subcellular localization by fractionation followed by RT-qPCR identified that PITAR is primarily located in the nucleus (**Figure 2A**). Similarly, the RNA In situ hybridization using RNAScope technology revealed that PITAR is located 80% in the nucleus (**Figure 2B**). PITAR silencing by two different siRNAs, # 1 and # 2 (**Figure 2C**), reduced cell proliferation, viable cell count, colony formation, arrested cells in G0/G1 phase, and induced apoptosis in U87 glioma cell line (**Figure 2D, E, F, G, and H).** PITAR silencing in U343 glioma cells also showed similar results (**Supplementary** Figure 2B**, C, D, E, and F**). Further, PITAR silencing showed more sensitivity to DNA-damaging agents, Adriamycin, and Temozolomide (**Figure 2I**). Interestingly, PITAR silencing did not affect the viable cell count in immortalized astrocytes (**Figure 2J)**. Overall, PITAR is a cancer-testis lncRNA that displays GBM and GSC-specific higher expression. We also demonstrate that PITAR promotes the growth of glioma cells and confers resistance to DNA-damaging agents.

**Figure 2:**
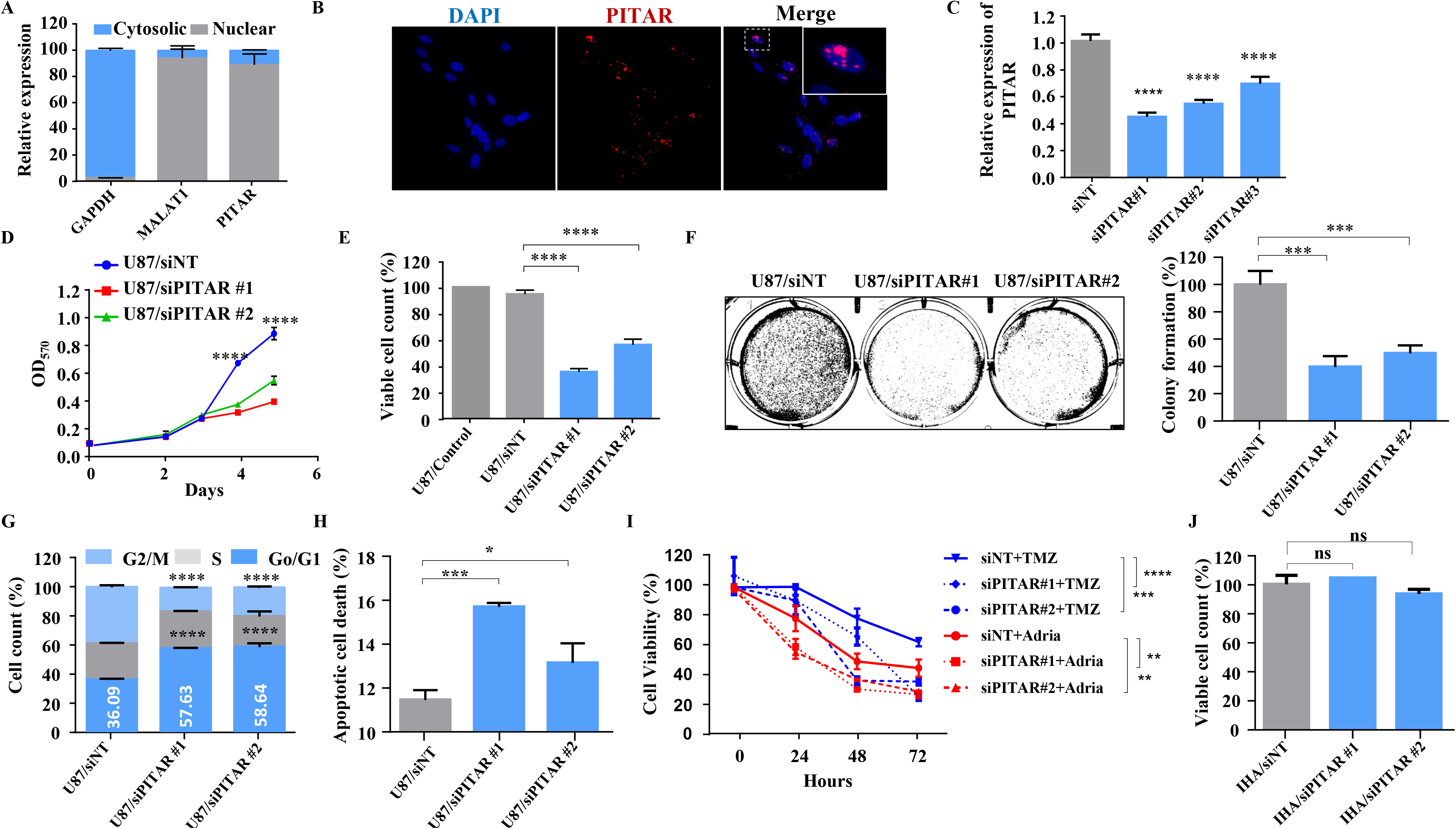
Glioblastoma cell proliferation and chemosensitivity altered by PITAR silencing. **A.** Plot depicts subcellular fractionation of U87 cells followed by quantification using qRT-PCR. MALAT1 is a positive control for nuclear gene expression, and GAPDH is a positive control for cytoplasmic gene expression. **B.** The RNAScope images of PITAR (red) and the DAPI (nucleus, blue) counterstain in U87 cells. **C.** The knockdown efficiency of siRNAs (siPITAR#1, siPITAR#2, and siPITAR#3) against PITAR was measured by qRT-PCR. **D.** The cell proliferation was measured by MTT assay upon PITAR knockdown in U87 cells. **E.** The viable cell count was measured using a viable cell counter following the Trypan blue method. **F.** Colony formation assay was performed upon PITAR knockdown compared to siNT in U87 cells. **G.** Cell cycle analysis was performed in PITAR-silenced U87 cells. **H.** The apoptotic cell death was measured by Annexin V/PI staining in PITAR-silenced U87 cells. **I.** The chemosensitivity upon PITAR silencing was measured by MTT assay against Adriamycin (0.25µg/ml) and Temozolomide (300µM) in U87 cells compared to control cells. **J.** The viable cell count of human astrocytes (IHA) was measured upon PITAR silencing. Data are shown as mean ± SD (n=3). \*\*\**P*-value <0.001, \*\**P*-value <0.01, \**P*-value <0.05.

### TRIM28 (Tripartite Motif Containing 28) mRNA is the direct target of PITAR

To identify PITAR interacting RNAs, we carried out an integrated analysis of RNA-Seq data from Chromatin Isolation by RNA Purification (ChIRP), performed using biotinylated PITAR-specific antisense probes and RNA-Seq data of PITAR silenced glioma cells. ChIRP was carried out using an odd (n=7) and even (n=7) set of biotinylated PITAR-specific antisense probes (**Supplementary** Figure 2A). RNA-Seq data of ChIRP RNA and further independent validation of ChIRP RNA with RT-qPCR showed an efficient pulldown of PITAR by both odd and even probes compared to LacZ (Control) probes (**Figure 3A and B**). The ChIRP data demonstrated that 827 mRNAs were enriched in the pulldowns using even and odd antisense probes compared to LacZ probes (**Supplementary Table 2)**. To choose the physiologically relevant target(s) for further studies, we intersected the ChIRP RNAs with i) GBM-associated differentially expressed transcripts that show a significant positive correlation (p <0.05 and r > 0.25) with PITAR transcript (**Supplementary Table 3)**, ii) GBM upregulated transcripts (**Supplementary Table 4)**, and iii) GSC upregulated transcripts (**Supplementary Table 5)**. This analysis identified 15 transcripts as potential targets of PITAR (**Figure 3C, D)**. Parallelly, RNA-Seq data of PITAR silenced U87 cells identified 946 differentially regulated genes (526 upregulated and 420 downregulated) (**Figure 3E; Supplementary Table 6)**. Gene ontology analysis of DEGs showed significant enrichment of biological processes such as “cell cycle,” “apoptosis,” and “p53” (**Figure 3F**). Furthermore, the Gene Set Enrichment Analysis (GSEA) showed significant enrichment of p53-regulated gene networks (**Supplementary Table 7**; **Figure 3G),** indicating that PITAR executes its functions by interfering with p53 functions.

**Figure 3:**
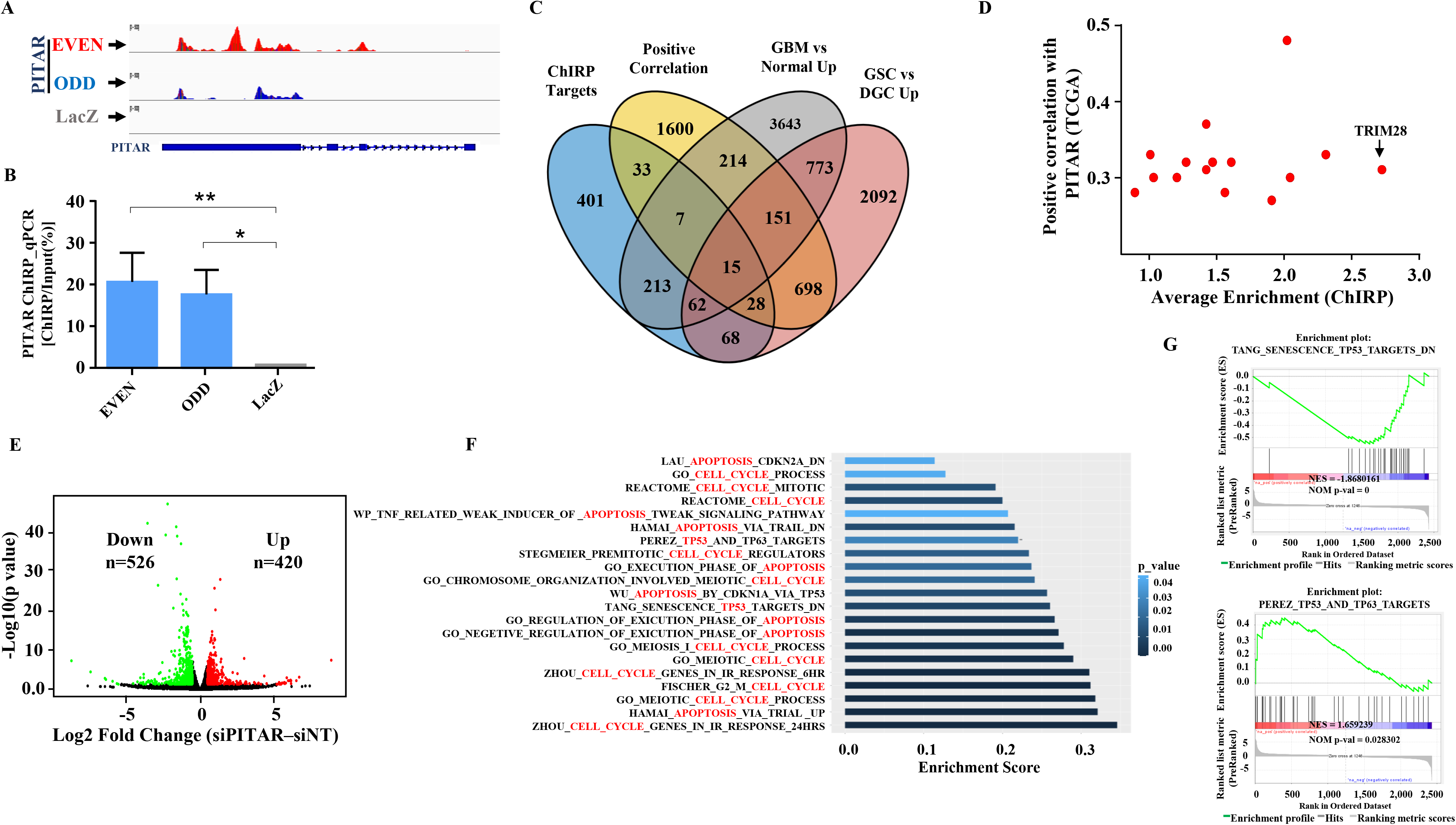
Identification of PITAR targets. **A.** Genomic track for PITAR derived from ChIRP-RNA sequencing using Odd, Even, and LacZ antisense probe. **B.** PITAR Pulldown by ChIRP assay was quantified using qRT-PCR. **C.** The Venn diagram represents the association of four datasets (ChIRP enriched genes, PITAR positive correlated genes from TCGA, GBM vs. normal upregulated genes from TCGA, and GSC vs. DGC upregulated genes from GSE54791). **D.** The selected fifteen genes from the Venn diagram are plotted in the scatter plot, and an arrow marked TRIM28 as a selected target. **E.** The volcano plot depicts up-regulated (n=420) and down-regulated (n=526) genes upon PITAR knockdown compared to siNT. The gene expression matrix between siPITAR and siNT was used to construct a volcano plot to visualize differentially expressed genes. **F.** Gene set enrichment analysis (GSEA) of differentially regulated genes was performed based on PITAR expression level at log2fold >0.58 and p<0.05. **G.** The GSEA plots depict the enrichment of p53 up and down target gene sets, results derived from PITAR-silenced U87 cells. Data are shown as mean ± SD (n=3). \*\*\**P*-value <0.001, \*\**P*-value <0.01, \**P*-value <0.05.

To explore the functional role of PITAR in p53-dependent gene expression, we chose TRIM28 among the ChIRP targets considering its role as p53-specific E3 ubiquitin ligase (Wang *et al*., 2005b). TRIM28, also known as KAP1 (Krüppel-Associated Box (KRAB)-Associated Protein 1), is a multidomain protein involved in various biological functions (Czerwińska, Mazurek and Wiznerowicz, 2017). TRIM28 inhibits p53 through HDAC1-mediated deacetylation and direct ubiquitination in an MDM2-dependent manner (Wang *et al*., 2005a). RNA-Seq data of the ChIRP assay showed a specific pulldown of TRIM28 mRNA by PITAR-specific odd and even antisense probes compared to LacZ probes (**Supplementary** Figure 3A). TRIM28 is significantly upregulated in multiple GBM cohorts (**Supplementary** Figure 4A**, B, C, D, and E**). Consistent with the ChIRP data, TRIM28 and PITAR transcripts showed a significant positive correlation in all GBM cohorts (**Supplementary** Figure 4F**, G, H, I, and J**). Like PITAR, the TRIM28 transcript is also expressed at higher levels in GSCs (**Supplementary** Figure 4 **K**), with a significant positive correlation between their expression in GSCs (**Supplementary** Figure 4 **L**). Further, TRIM28 protein showed higher levels in GBM as measured by quantitative proteomics data from the Clinical Proteomic Tumor Analysis Consortium (CPTAC) and immunohistochemistry from the Protein atlas (**Supplementary** Figure 4M and N).

Next, we investigated the specificity and impact of the interaction between PITAR and TRIM28 transcripts. Independent validation of ChIRP RNA with RT-qPCR showed an efficient pulldown of PITAR by PITAR-specific probes compared to LacZ (Control) probes (**Figure 4A**). Noncoding RNAs interact with target mRNAs through direct or indirect RNA-RNA interactions (Faghihi *et al*., 2008; Gong and Maquat, 2011; Kretz *et al*., 2012; Engreitz *et al*., 2014; Mahale *et al*., 2022a). Enrichment of TRIM28 mRNA in the PITAR ChIRP pull-down indicates the presence of potential TRIM28 mRNA: PITAR lncRNA interactions. Hence, we checked for the possible direct interaction between PITAR and TIRM28 by performing an RNA-RNA interaction analysis using the IntaRNA tool (http://rna.informatik.uni-freiburg.de/IntaRNA/Input.jsp) (Mann, Wright and Backofen, 2017; Gawronski *et al*., 2018). This analysis identified the most energetically favorable 80 bp interacting region between the 3’ UTR of the TRIM28 transcript and the first exon of PITAR (**Figure 4B; Supplementary Table 8)**. Next, we performed multiplexed RNAScope in U87 cells to evaluate the interaction between PITAR and TRIM28 in the cellular environment. The co-staining of U87 cells for TRIM28 transcript with the PITAR or PDE2A and PITAR with PDE2A demonstrated a specific colocalization of PITAR and TRIM28 transcripts, mainly in the nucleus. The extent of colocalization was much greater than that expected from coincidental colocalization with a similar abundant transcript, such as PDE2A (**Figure. 4C; Supplementary** Figure 3C).

**Figure 4:**
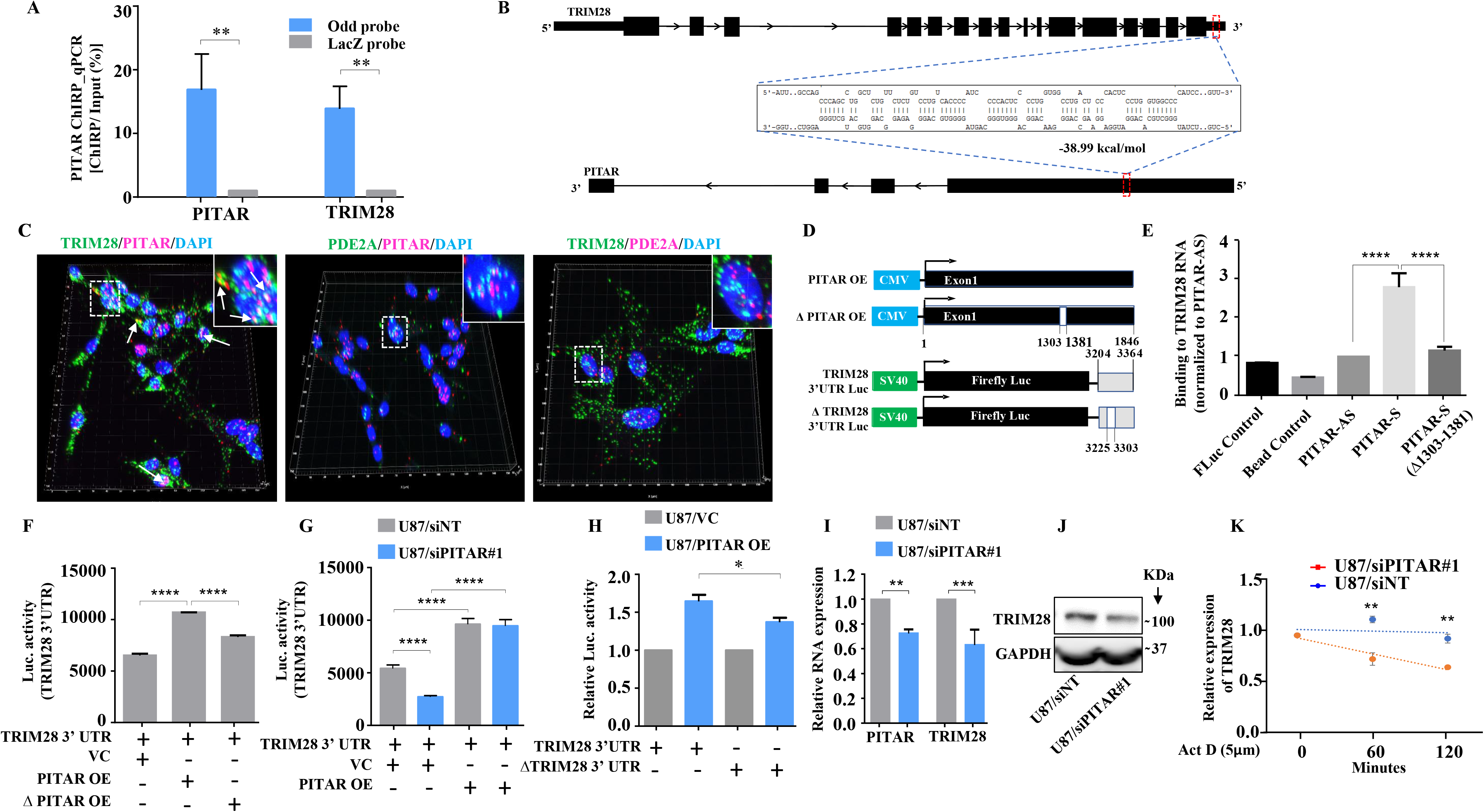
PITAR regulates the expression of TRIM28 by physical interaction with TRIM28. **A.** The qRT-PCR of PITAR and TRIM28 RNA was performed in ChIRP-RNA pull-down samples. The Probes for Odd and LacZ were used to pull down endogenous PITAR and interacting TRIM28 mRNA in U87 cells. **B.** Schematic represents the predicted RNA–RNA interaction between PITAR and the 3′ UTR of TRIM28. **C**. RNAScope images of co-localized signals of PITAR (red) and TRIM28 (green) in U87 cells. The panel shows the 3D reconstructed cell images of the merged 2D image (Imaris image analysis software). Yellow dots shown by white arrows depict co-localized PITAR (red) and TRIM28 (green). Magnified co-localized puncta were shown in the inset at the upper right corners, indicated by a white dotted box. The panel shows the RNAScope images of the localization of PITAR (red), TRIM28 (green), and PDE2A (green/red). PDE2A RNA was used as a negative control. The indicative scale bar on the images is 50µm. **D.** Schematic of vector plasmid construct of PITAR OE, ΔPITAR OE, TRIM28 3′ UTR, and ΔTRIM28 3′ UTR. **E.** PITAR interaction with the TRIM28 3′ UTR was measured using an in vitro RNA–RNA interaction assay and compared to a panel of control RNAs (PITAR antisense, Fluc control, Bead control). The binding affinity was quantified by qRT-PCR analysis of the TRIM28. Data were normalized to the PITAR-AS control. **F.** The Luc activity of TRIM28 3′ UTR was measured after the ectopic expression of PITAR and ΔPITAR in U87 cells using a luciferase reporter assay. **G.** Luciferase assay was performed in PITAR silenced U87 cells co-transfected with VC and PITAR OE vector. **H.** The Firefly luciferase activity was measured in U87 cells containing a deleted PITAR binding site of TRIM28 3’UTR (ΔTRIM28 3′ UTR), co-transfected with VC and PITAR OE. **I.** Relative expression of TRIM28 in PITAR-silenced U87 cells was measured by qRT-PCR. **J**. The TRIM28 protein expression was measured by immunoblotting. **K.** TRIM28 transcript was measured at indicated time points post Actinomycin D (5 μg/ml) treatment in siNT and siPITAR-transfected U87 cells by qRT-PCR. The log2 ratio of the remaining TRIM28 was plotted using linear regression after normalizing to the 0th hour of the respective condition. Data are shown as mean ± SD (n=3). \*\*\**P*-value <0.001, \*\**P*-value <0.01, \**P*-value <0.05.

To further confirm the interaction between PITAR and TRIM28, we cloned the first exon under the CMV promoter (PITAR OE) for exogenous overexpression and in vitro synthesis of biotin-labeled sense PITAR RNA (**Figure 4D**). First, we checked the ability of biotin-labeled PITAR sense RNA (corresponding to the first exon made from the PITAR OE construct) to bring down the TRIM28 mRNA from total RNA. PITAR-specific sense, but not antisense, biotinylated RNA brought down TRIM28 RNA efficiently (**Figure 4E, compare bar four with three**). PITAR sense RNA with a deletion of the 80 bp interacting region failed to bring down TRIM28 RNA (**Figure 4E, compare bar five with four**). The control biotin-labeled RNA (FLuc) and bead control did not bring down TRIM28 mRNA as expected. Further, we performed an antisense oligo-blocking experiment to validate the TRIM28 binding site on PITAR. Preincubation of fragmented total RNA with unlabelled PITAR antisense probe # 3, which is located close to the TRIM28 binding region on the exon 1 of PITAR, inhibited the ability of PITAR biotinylated antisense (odd) probe set to bring down TRIM28 (**Supplementary** Figure 3B). Next, we tested the ability of PITAR to regulate luciferase activity from the TRIM28-3’UTR-Luc construct (**Figure 4D**). The exogenous overexpression of PITAR (PITAR OE), but not PITAR with a deletion of 1303-1381 nucleotides (ΔPITAR OE), significantly increased the luciferase activity from the TRIM28-3’UTR-Luc construct (**Figure 4F, compare bars two and three with one**). In contrast, PITAR silencing significantly decreased the luciferase activity from TRIM28-3’UTR-Luc but was rescued by the exogenous overexpression of PITAR (**Figure 4G, compare bar two with one or four**). In contrast, the ability of PITAR exogenous overexpression to increase the luciferase activity from the ΔTRIM28-3’UTR-Luc construct, which has a deletion of 80 bp region corresponding to the PITAR binding region, is significantly reduced (**Figure 4H, compare bar four with two**). Next, to study the impact of PITAR interaction on TRIM28 expression, we measured TRIM28 transcript and protein levels in PITAR-silenced conditions. PITAR silencing significantly reduced TRIM28 mRNA (**Figure 4I, compare bar four with three**) and protein levels (**Figure 4I, compare lane two with one**). In actinomycin-treated U87 glioma cells, PITAR silencing reduced the TRIM28 mRNA half-life significantly compared with control cells (**Figure 4J, compare red line with blue line**). From these results, we conclude that PITAR interaction stabilizes TRIM28 mRNA to promote TRIM28 expression.

### PITAR inhibits p53 protein levels by its association with TRIM28 mRNA

TRIM28 inhibits p53 through HDAC1-mediated deacetylation and direct ubiquitination in an Mdm2-dependent manner (Wang *et al*., 2005a). We next investigated the impact of PITAR interaction with TRIM28 on p53. PITAR silencing in U87 glioma cells increased the luciferase activity from PG13-Luc, a p53-dependent reporter (El-Deiry *et al*., 1993) (**Supplementary** Figure 5A), decreased PITAR and TRIM28 mRNA levels, and increased CDKN1A mRNA levels with no change in p53 mRNA levels (**Figure 5A**). At the protein level, PITAR silencing decreased TRIM28 levels but increased p53 and CDKN1A levels (**Figure 5B, compare lanes two and three with one**). PITAR silencing in U343 cells also showed similar results (**Supplementary figure 5B and C**). In contrast, exogenous PITAR overexpression increased PITAR and TRIM28 mRNA levels and reduced CDKN1A mRNA levels without changing p53 mRNA levels (**Figure 5C**). Further, exogenous PITAR overexpression increased TRIM28 protein levels but decreased p53 and CDKN1A protein levels (**Figure 5D, compare lane two with one**), thus indicating that PITAR reduces p53 protein levels by regulating TRIM28. PITAR overexpression in U343 cells also showed similar results (**Supplementary** Figure 5D**, and E).** In cycloheximide-treated U87 glioma cells, the half-life of p53 increased (1.20 hr) under PITAR silenced conditions compared to control cells (0.50 hr) (**Figure 5E**). In good correlation, PITAR overexpression increased the ubiquitinated p53 levels in MG132-treated U87 glioma cells compared to the control condition (**Figure 5F, compare lane four with three**).

**Figure 5:**
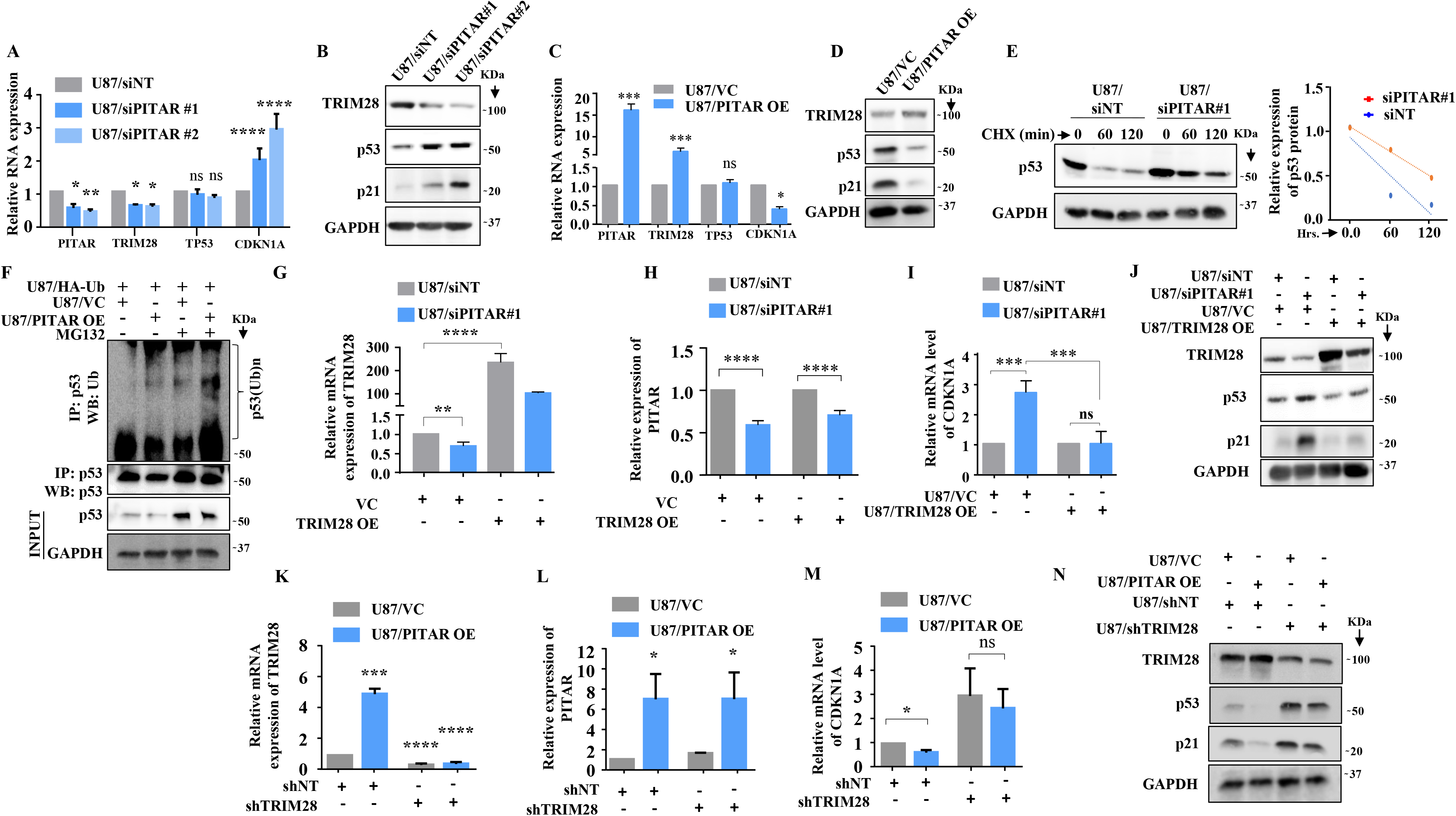
PITAR regulates wild-type p53 protein levels via TRIM28-mediated ubiquitination. **A.** Relative expression of PITAR, TRIM28, TP53, and CDKN1A was quantified by qRT-PCR in PITAR-silenced U87 cells compared to siNT. **B.** The protein expression of TRIM28, p53, and p21 was measured by immunoblotting in PITAR-silenced U87 cells compared to siNT. **C & D.** Cells were transfected with pcDNA3.1-PITAR (PITAR OE)/ empty vector control plasmid (pcDNA3.1) and harvested 48h post-transfection for qRT-PCR (PITAR, TRIM28, TP53, and CDKN1A) and immunoblotting with indicated antibodies (TRIM28, p53, and p21). GAPDH served as the control. **E.** The Half-life of the p53 protein was measured in PITAR-silencing (siPITAR) and control (siNT) U87 cells with the treatment of cycloheximide (CHX; 50 μg/mL). The relative expression of the remaining p53 was plotted using linear regression after normalizing to the 0th hour of the respective condition. **F.** The endogenous level of p53 ubiquitination was measured in pcDNA3.1-PITAR (PITAR OE)/empty vector plasmid (pcDNA3.1) stable U87 cells by p53 immunoprecipitation followed by immunoblotting with the indicated antibodies in the presence and absence of MG132. **G, H & I.** The relative expression of PITAR, TRIM28, and CDKN1A was measured by qRT-PCR in U87/siPITAR#1 and U87/siNT cells with exogenously overexpressed TRIM28 conditions. **J.** The protein expression of TRIM28, p53, and p21 was measured by immunoblotting in U87/siPITAR#1 and U87/siNT cells with exogenously overexpressed TRIM28 condition. **K, L & M.** The relative expression of PITAR, TRIM28, and CDKN1A was measured by qRT-PCR in U87/shTRIM28 and U87/shNT cells with exogenously overexpressed PITAR conditions. **N.** The protein expression of TRIM28, p53, and p21 was measured by immunoblotting in U87/shTRIM28 and U87/shNT cells with exogenously overexpressed PITAR conditions. Data are shown as mean ± SD (n=3). \*\*\**P*-value <0.001, \*\**P*-value <0.01, \**P*-value <0.05.

To confirm that TRIM28 mediates PITAR regulation of p53, we checked the ability of exogenously overexpressed TRIM28 (using a 3’UTRless TRIM28 construct) to rescue the phenotype in PITAR-silenced cells. TRIM28 overexpression resulted in a several-fold increase in TRIM28 mRNA levels in U87/siNT cells (**Figure 5G, compare bar three with one).** While this increase was significantly affected by PITAR silencing, the TRIM28 transcript levels remained high in U87/siPITAR cells (**Figure 5G, compare bar four with two).** As expected, TRIM28 overexpression did not affect the PITAR mRNA levels in U87/siNT and U87/siPITAR cells (**Figure 5H**). In TRIM28 overexpressing cells, PITAR silencing failed to increase luciferase activity from PG13-Luc (**Supplementary** Figure 5F**, compare bar four with three)** and CDKN1A transcript levels (**Figure 5I, compare bar four with three**). More importantly, PITAR silencing failed to increase p53 and CDKN1A protein levels in TRIM28 overexpressing U87 glioma cells (**Figure 5J, compare lane four with three**). Next, we tested the ability of PITAR overexpression to inhibit p53 functions in TRIM28 silenced cells. Exogenous overexpression of PITAR failed to increase the TRIM28 in TRIM28 silenced condition (**Figure 5K and L**). In addition, exogenous overexpression of PITAR failed to repress luciferase activity from PG13-Luc (**Supplementary** Figure 5G**, compare bar four with three)**, repress CDKN1A transcript levels (**Figure 5M, compare lane four with three**), and protein levels of p53 and CDKN1A (**Figure 5N, compare lane four with three**) in TIRM28 silenced cells. These results establish that PITAR inhibits p53 through its interaction with TRIM28 mRNA.

Next, we tested the requirement of WT p53 for the growth-promoting functions of PITAR as shown above (**Figure 2 and 5**). PITAR silencing inhibited colony formation efficiently in U87/shNT cells but not in U87/shp53#1 (**Supplementary** Figure 6A**, B, and C**). As shown before, PITAR silencing significantly increased CDKN1A and MDM2 transcript levels in U87/shNT cells but was compromised significantly in U87/shp53#1 (**Supplementary** Figure 6D). We also used glioma stem-like cells (GSCs) for this purpose. The ability of RG5, a patient-derived glioma stem cell line containing WT p53 (**data not shown**), to grow as a neurosphere was investigated in PITAR-silenced and overexpressed conditions. PITAR silencing inhibited the RG5 GSC growth in the neurosphere growth assay and limiting dilution assay (**Supplementary** Figure 7A**, B, C, and D**). PITAR silencing resulted in a decrease in PITAR and TRIM28 but an increase in CDKN1A transcript levels (**Supplementary** Figure 7E). As expected, there was no change in p53 transcript levels (**Supplementary** Figure 7E). The exogenous overexpression of PITAR promoted the neurosphere growth by RG5 (**Supplementary** Figure 7F**, G, and H**) and increased PITAR and TRIM28 transcript levels but decreased the CDKN1A transcript levels with no change in p53 transcript levels (**Supplementary** Figure 7I). In contrast, the growth of MGG8, a patient-derived glioma stem cell culture containing mutant p53 (**data not shown**), is not affected by PITAR silencing (**Supplementary** Figure 7J**, K, L, M, and N**). We conclude from these results that PITAR growth-promoting functions of glioma cells require wild-type p53.

### PITAR is induced by DNA damage in a p53-independent manner, which in turn diminishes the DNA damage response by p53

As p53 is activated by DNA damage (Shieh *et al*., 1997), we investigated the ability of PITAR to inhibit DNA damage-induced p53. Adriamycin-induced luciferase activity from PG13-Luc (**Supplementary** Figure 8A**, compare bar four with three**) and CDKN1A mRNA levels are significantly reduced by PITAR overexpression (**Figure 6A, compare bar four with three**). At the protein level, PITAR overexpression significantly reduced adriamycin-induced total p53, acetylated p53, and CDKN1A levels (**Figure 6B, compare lane four with three**). In addition, the viable cell count increased significantly in adriamycin-treated cells upon PITAR overexpression (**Figure 6C, compare bar four with three**). Similarly, the extent of G2/M arrest seen in adriamycin-treated cells is reduced significantly (18.34 %) upon PITAR overexpression (**Figure 6D; 45.80 % is reduced to 37.40%**).

**Figure 6:**
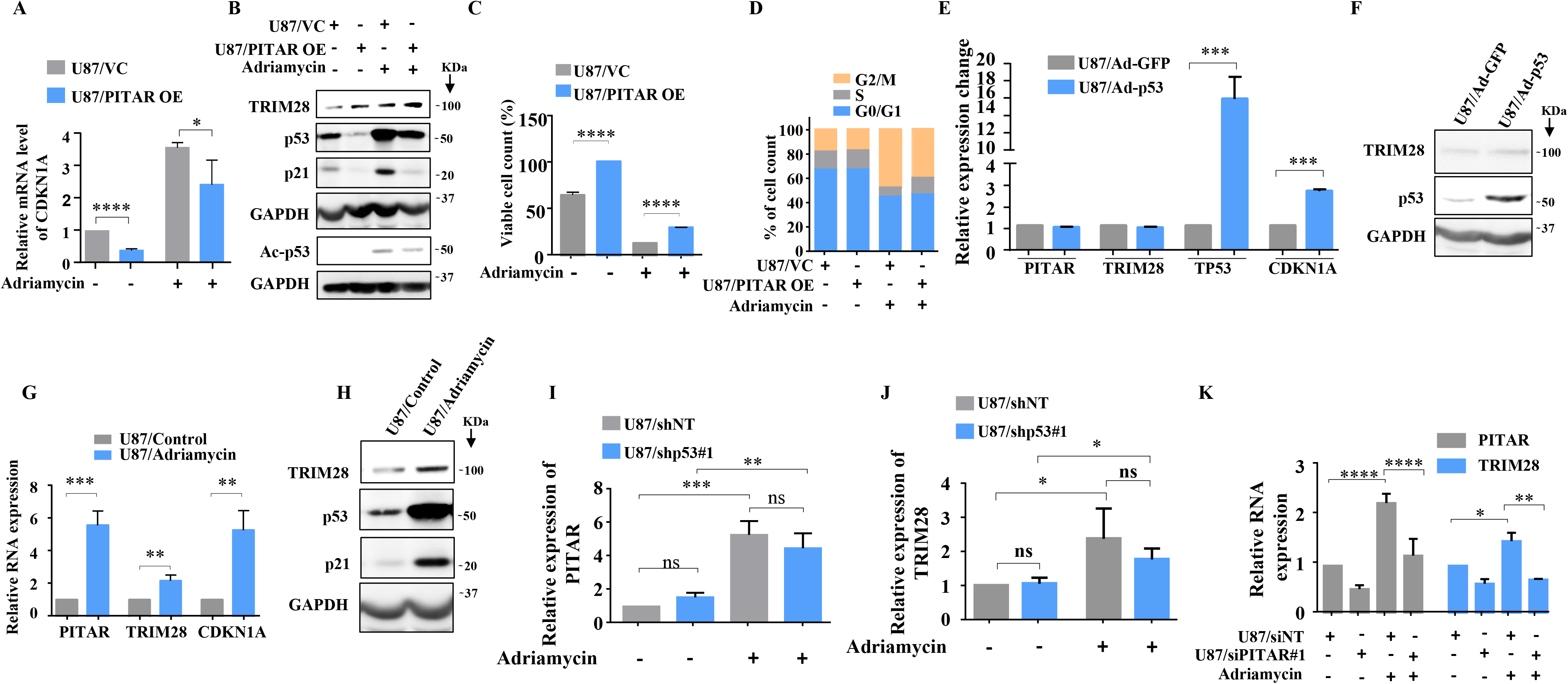
DNA damage-induced PITAR diminishes the DNA damage response by p53 through TRIM28. **A.** The relative expression of CDKN1A was measured in the presence and absence of Adriamycin by qRT-PCR in PITAR OE/VC U87 cells. **B.** The protein expression of TRIM28, p53, ac-p53, and CDKN1A was measured in the presence and absence of Adriamycin by immunoblotting in PITAR OE/VC U87 cells. GAPDH served as the control. **C.** The viable cell count was performed in the presence and absence of Adriamycin in PITAR OE/VC U87 cells. **D.** The Cell cycle analysis was performed in the presence and absence of Adriamycin in PITAR OE/VC U87 cells. **E.** The relative expression of PITAR, TRIM28, TP53, and CDKN1A was measured by qRT-PCR in Ad-p53 and Ad-GFP-infected U87 cells. **F.** The protein expression of TRIM28 and p53 was measured by immunoblotting in Ad-p53 and Ad-GFP-infected U87 cells. **G.** The relative expression of PITAR, TRIM28, and CDKN1A was measured in the presence and absence of Adriamycin by qRT-PCR in U87 cells. **H.** The immunoblot depicting the expression of TRIM28, p53, and p21 upon treatment of Adriamycin. **I & J.** The qRT-PCR was performed to measure the relative expression of PITAR and TRIM28 in the p53 knockdown condition upon Adriamycin treatment. **K.** The relative expression of PITAR and TRIM28 was measured by qRT-PCR in Adriamycin-treated PITAR-silenced U87 cells. Data are shown as mean ± SD (n=3). \*\*\**P*-value <0.001, \*\**P*-value <0.01, \**P*-value <0.05.

To verify the possible existence of a negative feedback loop between PITAR and p53, we checked the ability of p53, expressed from a recombinant adenovirus, to induce PITAR transcript levels. We found that Ad-p53 induced p53 and CDKN1A mRNA and p53 protein levels compared to the control virus (Ad-GFP) but did not alter the PITAR and TRIM28 transcript levels (**Figure 6E and F**), thus confirming that neither PITAR nor TRIM28 is the direct target of p53. Interestingly, DNA damage by adriamycin treatment also induced PITAR, TRIM28 transcript, and TRIM28 protein levels besides CDKN1A transcript and protein levels of p53 and CDKN1A (**Figure 6G and H**). Our results also show that adriamycin treatment induced PITAR and TRIM28 transcript levels efficiently in p53 silenced cells (**Figure 6I and J, compare bar four with three),** indicating DNA damage-mediated PITAR induction is p53 independent. Further, PITAR and TRIM28 induction by DNA damage is dependent on ATM/ATR kinase as the pretreatment of glioma cells with CGK733, a small molecule inhibitor of ATM/ATR kinase, prevented the adriamycin-mediated induction of PITAR and TRIM28 transcript levels (**Supplementary** Figure 8B) and TRIM28 protein level (**Supplementary** Figure 8C) thus reiterating the importance of ATM/ATR kinases, as previously demonstrated (Tibbetts et al., 1999; Cheng and Chen, 2010), in DNA damage response pathway. Interestingly, the DNA damage-induced TRIM28 transcript upregulation is found to be dependent on PITAR upregulation, as adriamycin failed to induce TRIM28 in PITAR-silenced U87 cells (**Figure 6K, compare bar eight with seven**). This was further confirmed by the fact that adriamycin-induced luciferase activity from TRIM28 3’-UTR-Luc is inhibited by CGK733 treatment (**Supplementary figure 8D, compare bar three with two**), suggesting the requirement of PITAR for the TRIM28 transcript upregulation in DNA-damaged cells. These results establish that PITAR is DNA damage-inducible in a p53-independent manner, and it inactivates p53 through its association with TRIM28 mRNA.

The p53-Mdm2 autoregulatory negative feedback loop controls the extent and duration of p53 response upon DNA damage (Wu *et al*., 1993; Haupt *et al*., 1997; Kubbutat, Jones and Vousden, 1997; Zhang *et al*., 2009). Since TRIM28 association with Mdm2 contributes to p53 inactivation (Wang *et al*., 2005b; Czerwińska, Mazurek and Wiznerowicz, 2017), we hypothesized that PITAR, through its association with TRIM28 may also contribute to the control of DNA damage response by p53. We tested this possibility by measuring the p53 response in PITAR-silenced cells. U87/siNT and U87/siPITAR#1 cells were exposed to 7 Gy of ionizing radiation, and steady-state levels of p53, Mdm2, and p21 proteins were determined at hourly intervals. In U87/siNT cells, p53 protein peaked around 3 hrs of irradiation, while mdm2 protein peaked at 7 hrs of irradiation, as expected (**Supplementary** Figure 8E**, F, and G; blue line**). p21 protein largely followed similar kinetics to that of the Mdm2 protein (**Supplementary** Figure 8E and H**; blue line**). In U87/siPITAR#1 cells, while the overall kinetics of p53, Mdm2, and p21 protein were found to be similar in terms of duration, the extent of p53 activation was much stronger (**Supplementary** Figure 8E**, F, G, and H; red line).** The levels of mdm2 and p21 proteins followed a similarly strong response in PITAR-silenced cells (**Supplementary** Figure 8E**, F, G, and H; red line).** From these results and our above findings, we conclude that DNA damage-induced PITAR diminishes the p53 response to DNA damage.

### PITAR promotes glioma tumor growth in a TRIM28-dependent manner and resistance to Temozolomide chemotherapy

To investigate the role of PITAR on tumor growth, we checked the ability of PITAR-silenced U87 glioma cells to form a tumor in an intracranial orthotopic tumor model using NIH nu/nu mice. We found that PITAR silencing (U87/siPITAR) significantly reduced tumor growth (**Figure 7A, B, and C; compare blue line with red line**) and enhanced mouse survival (**Figure 7D**). U87/siPITAR tumors showed reduced TRIM28 staining (**Figure 7E**). Next, we investigated the impact of PITAR overexpression on glioma tumor growth and Temozolomide (TMZ) chemotherapy. U87/PITAR OE cells-initiated tumors grew much faster than U87/VC cells (**Figure 7F, and G; compare red line with blue line; Supplementary** Figure 9A), thus confirming that PITAR overexpression promotes tumor growth. While the growth of the tumors formed by U87/VC glioma cells is inhibited substantially by TMZ chemotherapy, U87/PITAR OE tumors showed resistance to TMZ chemotherapy (**Figure 7F and G; compare pink line with green line; Supplementary** Figure 9A). U87/PITAR OE tumors showed higher TRIM28 and Ki67 (proliferation marker) staining but reduced p21 staining (**Supplementary** Figure 9B).

**Figure 7:**
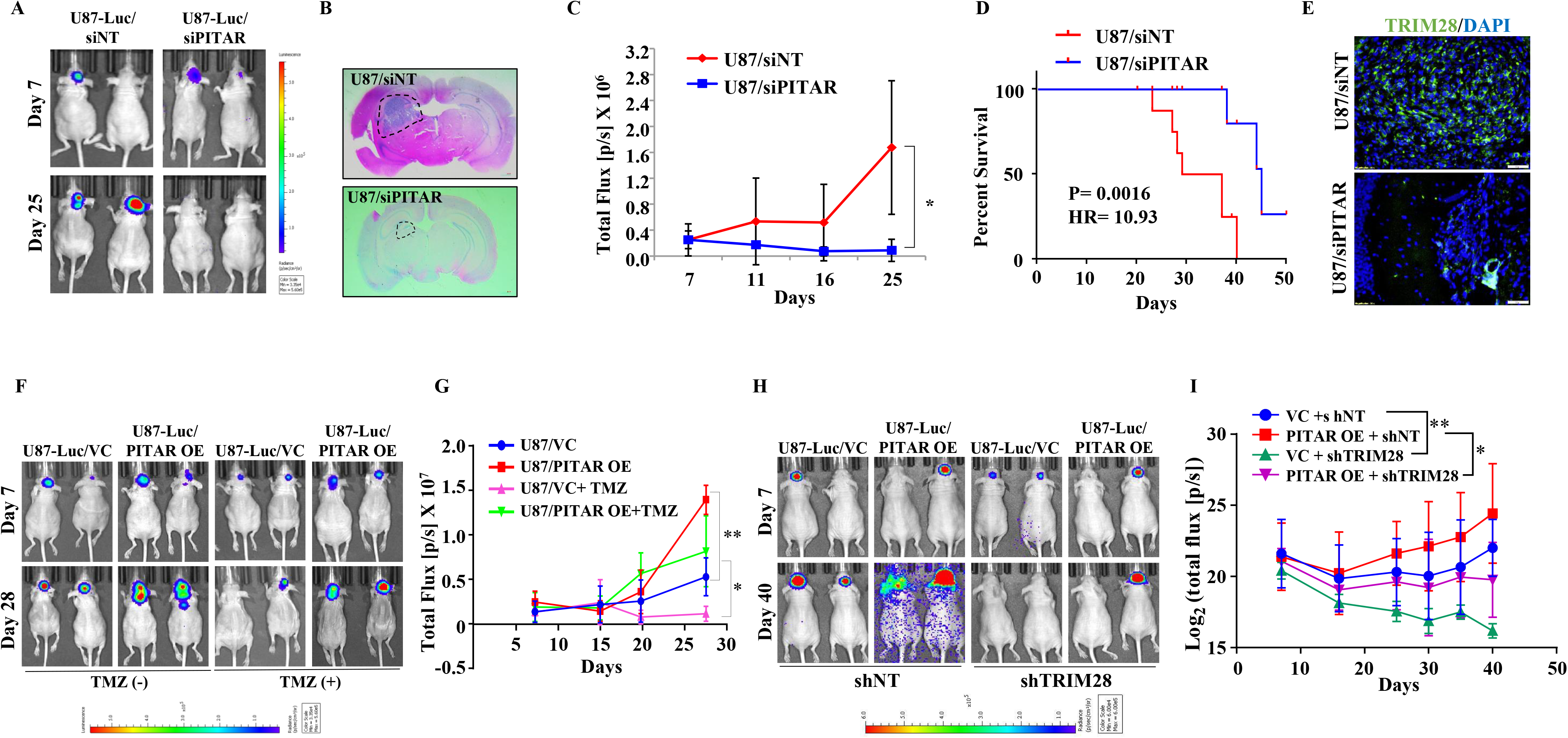
PITAR promotes glioma tumor growth and resistance to Temozolomide chemotherapy. **A.** Mice (NIH nu/nu) were injected with siNT/siPITAR#1 transfected U87-Luc cells (0.3x10^6^ cells/mice), and tumors were allowed to grow for 50 days, and the luminescence imaging was performed using the IVIS instrument (Perkin Elmer IVIS system). **B.** H&E staining was performed in formalin-fixed tumor-bearing (siNT and siPITAR#1) mouse brain sections. **C.** The tumor growth curve of siNT and siPITAR#1 tumor-bearing mice was quantified over time using IVIS. **D.** The Kaplan–Meier graph shows the survival of mice-bearing tumors formed by siNT and siPITAR#1 cells. **E.** The Immunohistochemistry assay was performed to show the TRIM28 protein expression in the tumor tissue section derived from U87-Luc/siNT and U87-Luc/siPITAR#1 tumors. The Green color represents the TRIM28 protein, and the blue depicts the nucleus stained with DAPI. Scale bar = 100 μm. **F.** Mice (NIH nu/nu) were injected with U87-Luc/PITAR OE and U87-Luc/VC cells (0.3x10^6^ cells/mice), and tumors were allowed to grow for 30 days. The tumor-bearing mice were treated with 100 mg/kg TMZ in 25% DMSO saline solution after 11 days by intraperitoneal injection for one week, and the luminescence imaging was performed using the IVIS instrument. **G.** The tumor growth curve of VC, PITAR OE, VC+TMZ, and PITAR OE+TMZ tumor-bearing mice was quantified over time using IVIS. **H.** Mice (NIH nu/nu) were injected with U87-Luc/PITAR OE+shNT, U87-Luc/VC+shNT, U87-Luc/PITAR OE+shTRIM28 and U87-Luc/VC+shTRIM28 cells (0.3x10^6^ cells/mice), and tumors were allowed to grow for 50 days. **I.** The tumor growth curve of VC+shNT, PITAR OE+shNT, VC+shTRIM28, and PITAR OE+shTRIM28 tumor-bearing mice was quantified over time using IVIS. Luminescence was evaluated twice per 10 days and before sacrifice. Bars indicate standard error. Data are shown as mean ± SD (n=3). \*\*\**P*-value <0.001, \*\**P*-value <0.01, \**P*-value <0.05.

To check the importance of TRIM28 in the glioma tumor growth-promoting functions of PITAR, we tested the ability of exogenously expressed PITAR to promote tumor growth in TRIM28-silenced cells. U87/PITAR OE/shNT cells formed a larger tumor compared to U87/VC/shNT cells (**Figure 7H and I; compare red line with blue line**). However, U87/PITAR OE/shTRIM28 glioma cells formed significantly smaller tumors compared to U87/PITAR OE/shNT cells (**Figure 7H and I; compare pink line with red line**). As expected, small tumors formed by U87/PITAR OE/shTRIM28 glioma cells showed reduced TRIM28 and enhanced p21 staining compared to large tumors formed by U87/PITAR OE/shNT glioma cells (**Supplementary** Figure 9C). These results demonstrate that PITAR promotes glioma tumor growth in a TRIM28-dependent manner and confers resistance to TMZ chemotherapy.

To further explore the clinical relevance of our findings, we investigated the survival significance of PITAR using the GBM transcriptome datasets. While PITAR and TRIM28 transcripts showed a significant positive correlation in the GBM full cohort and p53 wild-type cohort, there was no significant correlation in the p53 mutant cohort (**Supplementary** Figure 9D). However, survival analysis by univariate Cox regression revealed that PITAR transcript level predicted survival only in the p53 wild-type GBM cohort but neither in the full nor p53 mutant cohort (**Supplementary** Figure 9E). The PITAR transcript level was dichotomized on further analysis to elucidate the cut-off in predicting the prognosis. We found that high PITAR transcript levels predicted poor prognosis significantly compared to low PITAR transcript levels in the p53 wild-type cohort (**Supplementary** Figure 9G). However, the PITAR transcript levels failed to predict survival in full or p53 mutant GBM cohorts (**Supplementary** Figure 9F and H). These results prove that PITAR promotes tumor growth and therapeutic resistance by inactivating p53 by its association with TRIM28.

## Discussion

Research into cancer for many decades focussed on protein-coding genes. However, recent evolution in RNA-Seq technologies and bioinformatic methods to analyze transcriptome and genome has changed our perception of noncoding RNA (ncRNA) from “junk RNAs” to functional regulatory molecules that control various biological processes, such as chromatin remodeling, gene regulation at transcription, post-transcription, and post-translation level, post-translational modifications of proteins, and signal transduction pathways. It appears lncRNAs can influence various macromolecules such as DNA, RNA, and protein to execute specific biological responses and cell fate. Concerning cancer, lncRNAs are critical regulators of cancer and have been shown to influence cancer origin and progression by acting as tumor drivers and/or suppressors in a cancer-specific fashion (Anastasiadou et al., 2018; Slack & Chinnaiyan, 2019; Yan & Bu, 2021). Besides, several lncRNAs are identified as novel biomarkers and potential therapeutic targets for cancer (Qian, Shi and Luo, 2020). p53 inactivation is an essential step in cancer development and is associated with therapy resistance. While genetic alteration forms the major mode of p53 inactivation, alternate ways of p53 inactivation, such as amplification of MDM2 and deletion of p19arf, have been reported (Vogelstein, Lane and Levine, 2000; Olivier *et al*., 2002; Vousden and Lu, 2002; Vousden and Lane, 2007). Deregulated lncRNAs that modulate p53 activity positively and negatively have been reported (Jain, 2020). In this study, we discovered that PITAR, an oncogenic lncRNA, inhibits p53 by promoting its ubiquitination through its association with the mRNA of TRIM28 that encodes p53-specific E3 ligase. PITAR is activated by DNA damage in a p53-independent manner suggesting an incoherent feedforward loop.

Several deregulated lncRNAs acting as oncogenes and tumor suppressors in glioma and glioma stem cells have been identified (Paul *et al*., 2018; Wang and He, 2019; Yadav *et al*., 2021). In this study, we identify PITAR as a highly expressed lncRNA in GBM and GSCs and display oncogenic properties. PITAR is primarily located in the nucleus. Its downregulation by RNA interference inhibited glioma cell growth, and colony formation, arrested cells in the G1 phase, induced cell death, and sensitized cells to chemotherapy. An integrated analysis of PITAR-bound targets identified by ChIRP assay and differentially regulated genes in PITAR silenced cells identified PITAR as an inhibitor p53 through its association with the transcript encoding TRIM28.

TRIM28 (KAP1), an E3 ligase, is an MDM2 interacting protein, and it inhibits p53 through HDAC-mediated deacetylation and direct ubiquitination (Czerwińska et al., 2017; C. Wang et al., 2005). Our results show TRIM28 transcript is upregulated in GBM, as reported earlier (Czerwinska *et al*., 2017; Su, Li and Gao, 2018), and GSCs in multiple datasets. TRIM28 protein was also found to be high in GBM tissue samples. In addition, we found a significant positive correlation between PITAR and TRIM28, suggesting that PITAR expression is likely to promote TRIM28 expression. Our results show that PITAR-dependent TRIM28 expression regulation occurs at the post-transcription level via RNA: RNA interactions, where PITAR interaction with TRIM28 mRNA increases its stability. This interaction between PITAR and TRIM28 was confirmed by colocalization using a multiplexed RNAscope. We further demonstrated that the first exon of PITAR interacts with 3’ UTR of TRIM28. Thus, we have comprehensively established that high levels of PITAR in GBM promote TRIM28 expression by binding and stabilizing TRIM28 mRNA. TRIM28 inhibits p53 through its association with MDM2 (Wang *et al*., 2005a). Since our results show that PITAR binding to TRIM28 transcript promotes TRIM28 expression, PITAR is expected to inhibit p53. Indeed, our results show that PITAR inhibits both endogenous and DNA damage-induced p53 at the protein level without altering its transcript levels. Increased TRIM28 protein in high PITAR cells increased p53 ubiquitination and reduced p53 acetylation. Thus, we establish PITAR as a bona fide inhibitor of p53 through its association with TRIM28 mRNA.

Several cellular stresses, including DNA damage, hypoxia, and nucleotide deprivation, activate p53. Several lncRNAs have been shown to regulate p53 (Jain, 2020). LncRNAs such as LOCC572558, NBAT1, MT1JP, PSTAR, and MEG3 are shown to activate p53 using a variety of mechanisms (L. Liu *et al*., 2016; Zhu *et al*., 2016; Uroda *et al*., 2019; Qin *et al*., 2020; Mitra *et al*., 2021). In contrast, lncRNAs such as PURPL, MALAT1, H19, and RMRP inhibit p53 (C. Liu *et al*., 2016; Chen *et al*., 2017, 2021; Li *et al*., 2017). While PITAR is found to inhibit p53 through its interaction with TRIM28 mRNA, we also found that PITAR level is induced by DNA damage in a p53-independent manner. We show further that DNA damage induced PITAR inhibits the p53 response to DNA damage through its interaction with TRIM28 mRNA. Thus, based on our data, we propose a model wherein DNA damage not only activates p53, but it also activates PITAR, with oncogenic properties, to inhibit p53-dependent functions, thus creating an incoherent feedforward mechanism to attenuate the response of p53 to DNA damage and promote oncogenesis and therapy resistance in GBM (**Figure 8**).

**Figure 8:**
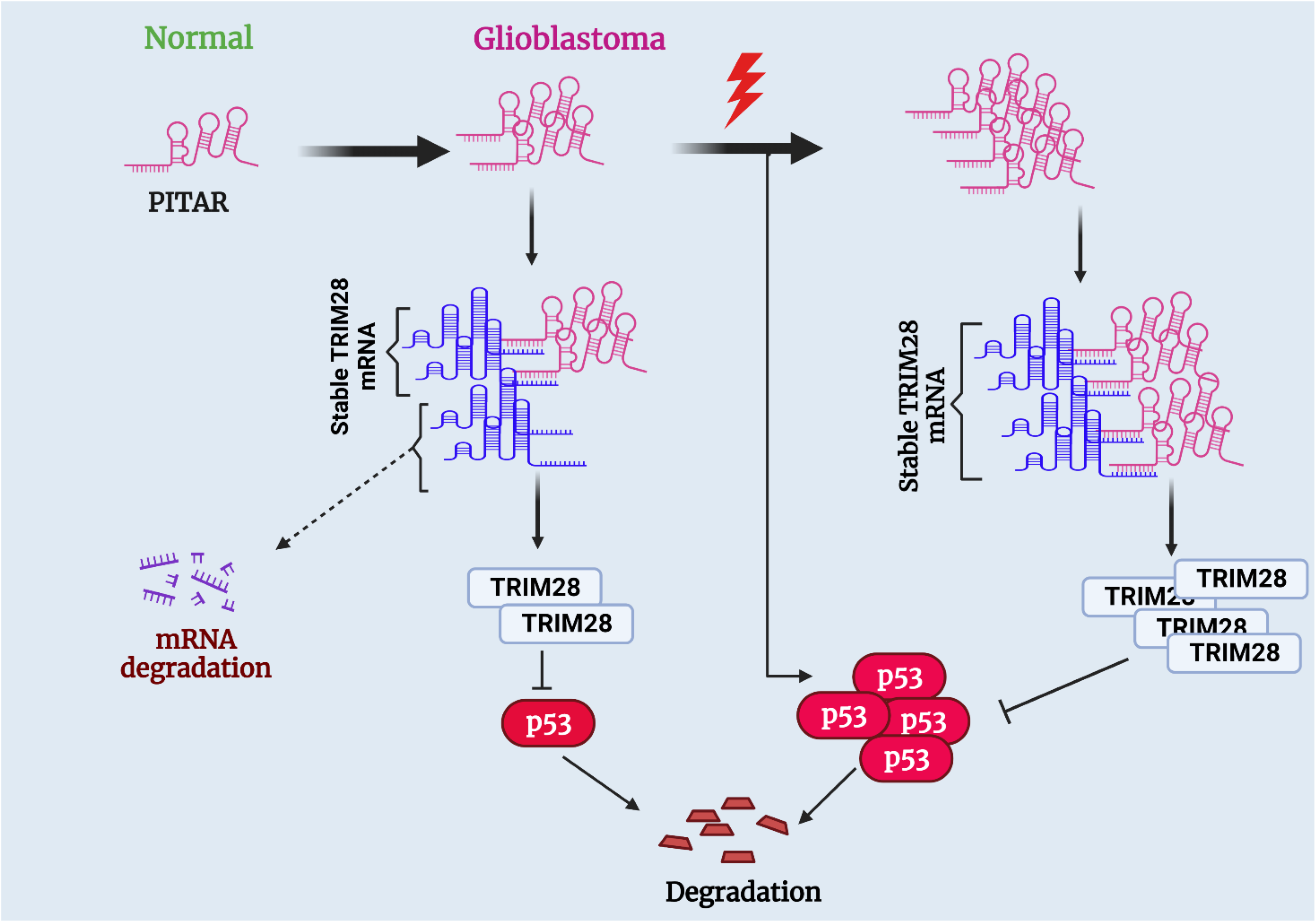
Proposed working model of this study. PITAR inhibits p53 by binding and stabilizing TRIM28 mRNA (created with BioRender.com).

Several oncogenic lncRNAs have been demonstrated as potential therapeutic targets in many cancer (Anastasiadou, Jacob and Slack, 2018; Slack and Chinnaiyan, 2019b; Yan and Bu, 2021). In GBM, glioma stem-like cells (GSCs) are the main culprits behind highly aggressive GBM progression (Visvanathan *et al*., 2018). It has been shown that GSCs alone can initiate glioma tumors in mouse models (Suvà *et al*., 2014). Cancer/testis (CT) antigens are a family of tumor-associated antigens expressed only in tumors but not in normal tissues except for the testis and placenta. Their tumor-specific expression and strong in vivo immunogenicity made them suitable for tumor-specific immunotherapy (Fratta *et al*., 2011; Gjerstorff, Andersen and Ditzel, 2015). Because of its high expression in GBM and testis normal tissue, PITAR is identified as CT lncRNA. PITAR silencing efficiently inhibited the growth of glioma cells and GSCs containing WT p53. Intracranial orthotopic mouse model experiments show that PITAR promotes glioma tumor growth and confers resistance to temozolomide chemotherapy. Our study establishes PITAR as an oncogenic CT lncRNA, which promotes oncogenesis by inactivating p53 by interacting with TRIM28 mRNA. Thus it can serve as a potential therapeutic target for GBM.

## Materials and Methods

### Tumor samples and clinical details

Tumor samples were collected from patients operating at the hospital (NIMHANS, Bangalore). A portion of the anterior temporal cortex resected during surgery for drug-resistant epilepsy patients served as a control brain sample (NIMHANS, Bangalore). We used GBM (n=79) and Normal (n=3) tissue to quantify PITAR and TRIM28 expression and correlation. The neuropathologist confirmed the histology as GBM, IDH wild type.

### Cell lines

Primary human tumor-derived GSCs Neurosphere (MGG4, MGG6, MGG8) are obtained from Wakimoto’s Lab, RG5 and 120-53 GSCs Neurosphere obtained from Sánchez-Gómez’s Lab, and other GSCs Neurosphere (GB1, GB2, GB3) are developed in our Lab. The Neurospheres were grown in Neurobasal medium (#21103049, Gibco) supplemented with l-glutamine, heparin, B27 supplement, N2 supplement, rhEGF, rhFGF-basic, penicillin, and streptomycin. To make single-cell suspensions for re-plating, the spheres were chemically dissociated after seven days of plating, using the NeuroCult Chemical Dissociation Kit (mouse, #05707) from Stem Cell Technologies, according to the manufacturer’s instructions. The monolayer culture GBM cell lines U87, U343, U252, LN229, A172, U373, and immortalized human astrocyte cell line IHA (NHA-hTERT-E6/E7) was obtained from Dr. Russell Pieper’s laboratory, University of California, San Francisco (San Francisco, CA), and cells are maintained in specific culture media with 5% CO2 and humidified incubator at 37 °C. The mycoplasma contamination was tested using RT–PCR. All cell lines are verified to be mycoplasma-free by EZdetect PCR Kit for Mycoplasma Detection (HiMedia).

### Plasmids

The shRNA plasmid (pLKO.1) for TP53 (TRCN0000003754) and TRIM28 (TRCN0000017998) were obtained from the TRC library (Sigma, IISc). The TRIM28 3’ UTR Luc vector plasmid was purchased from Origen, USA (# SC202088). PG13-Luc vector plasmid was obtained from Addgene, USA (#16442). The Flag-TRIM28 overexpression plasmid was bought from Addgene, USA (# 124960). The pRK5-HA-Ubiqitin-K48 plasmid was procured from Dr. Sashank Tripathi’s laboratory (Addgene_17605).

### Antibodies and reagents

Primary antibodies were purchased from the following commercial vendors: p53 (DO-1) from Santacruz; acetylated p53 (K382), p21, TRIM28, and Anti-Ub from Cell Signaling Technology; GAPDH and β-actin from Sigma Aldrich. Goat anti-mouse HRP conjugate (Bio-Rad #170-5047, WB 1:5000), goat anti-rabbit (H+L) secondary HRP conjugate (Invitrogen, #31460, WB 1:5000), goat anti-mouse IgG (H+L) highly cross-adsorbed secondary antibody, Alexa Fluor 488 (Invitrogen, #A-11029), goat anti-rabbit IgG (H+L) highly cross-adsorbed secondary antibody, Alexa Fluor 488 (Invitrogen, Cat# A-11034), goat anti-mouse IgG (H+L) highly cross-adsorbed secondary antibody, Alexa Fluor 594 (Invitrogen, Cat# A-11032), goat anti-rabbit IgG (H+L) highly cross-adsorbed secondary antibody, Alexa Fluor 594 (Invitrogen, Cat#A-11037). All Alexa Fluor conjugated antibodies were used at a dilution of 1:500 for IHC and ICC. The PCR primers were purchased from Sigma, and the Biotin-TEG DNA antisense oligos were designed in *Stellaris probe* designer and purchased from IDT. PITAR siRNAs and negative control oligos were purchased from Eurofins. Doxorubicin, Temozolomide, CGK733, MG132, Actinomycin D, and Cycloheximide were purchased from Sigma.

### Plasmid construction and RNA interference

The PITAR OE partial clone (∼1846 bp) was made in the pCDNA3.1 vector backbone, and the deletion clone (ΔPITAR OE) was made using a Q5 site-directed mutagenesis assay kit (#E0554S). The deletion clone of TRIM28 3’UTR (ΔTRIM28 3’UTR) was made using a Q5 site-directed mutagenesis assay kit. The PITAR Knockdown was performed using PITAR siRNA and transfected using Dharmafect1 transfection reagent. Cells were harvested at the 24th, 48th, 72nd, and 96th hr post-transfection to check for knockdown of desired genes at mRNA level by qRT-PCR.

To prepare the Lentivirus for shRNA (pLKO.1 vector), HEK-293T cells were transfected with shRNA plasmid and helper plasmids psPAX2 and pMD2.G using Lipofectamine 2000 (Invitrogen #11668027) in Opti-MEM (Invitrogen #22600-050) medium. After 6 hours of transfection, the Opti-MEM medium was replaced by a fresh DMEM supplemented with 10% FBS, and the virus was collected after 60 hours of transfection. The knockdown of p53 and TRIM28 was performed using shTP53 and shTRIM28 lentivirus infection, followed by puromycin selection. The exogenous expression of p53 was executed by using a recombinant adenovirus for p53 and compared to the control virus (Ad-GFP) which is amplified in HEK-293 cells.

### The identification of differentially expressed lncRNA and mRNA in GBM vs. control brain tissue and GSC vs. DGC

The Raw RNA sequencing data was obtained for GBM samples from TCGA (https://tcga-data.nci.nih.gov/tcga/). The whole RNA sequencing data were aligned using the PRADA tool (doi: 10.1093/bioinformatics/btu169). Duplicate removal was carried out using Picard 1.73 (http://broadinstitute.github.io/picard/), and the lncRNAs were annotated as per the Gencode Version 19 annotation file (http://www.gencodegenes.org/releases/19.html). The RNA-seq data for GSC vs. DGC was obtained from GSE54791 (Suva et al. 2014). The gene expression matrix obtained was log2 transformed. Further, the count expression matrix for GBM and Normal was obtained from TCGA, CPM was normalized using library edgeR, and limma using R. Common protein-coding genes and lncRNA were chosen, and a scatterplot was constructed among them. The pan-cancer data for the gene expression profile was derived from the TCGA browser (https://tools.altiusinstitute.org/tcga/?gene=FAM95B1). The tissue-specific gene expression and regulation were analyzed using the GTEx portal (Samples were collected from 54 non-diseased tissue sites across nearly 1000 individuals). The quantitative proteomics data of tumors was derived from the CPTAC data portal.

### RNA-sequencing

Total RNA was extracted from siNT and siPITAR cells using the Trizol method (Qiagen). RNA quality was assessed using an Agilent TapeStation system, and a cDNA library was made. According to published protocols, each sample was sequenced using the Illumina HiSeq 2000 (with a 100-nt read length). To quantitate the abundance under this condition. The raw reads were quality-analyzed, and upon satisfactory assessment, Kallisto was used with default parameters to quantitate and obtain TPM-normalized gene abundance. The fold change was brought using Deseq2. IGV was used to visualize the raw reads. David and GSEA were performed both at log2fold 0.58 and p<0.05. The gene expression matrix between siPITAR and Control was used to construct a volcano plot to visualize differentially expressed genes.

### Gene set enrichment analysis (GSEA)

The differentially expressed genes between the siNT and siPITAR (as identified from RNA-seq) were pre-ranked based on fold change and used as an input to perform GSEA. All the gene sets available in the Molecular Signature Database (MSigDB, roughly 18,000 gene sets) were used to run the GSEA. We filtered out the cell cycle, apoptosis, and p53 pathway-related gene sets to identify that most of them were significantly enriched in the siPITAR over the siNT. We acknowledge using the GSEA software and MSigDB (http://www.broad.mit.edu/gsea/) (Subramanian *et al*., 2005).

### RNA isolation and real-time Quantitative RT-PCR analysis

Total RNA was isolated using TRI reagent (Sigma, U.S.A.), and 2µg of RNA was reverse transcribed with the High-capacity cDNA reverse transcription kit (Life Technologies, USA) according to the manufacturer’s protocol. qRT-PCR was performed using DyNAmo ColorFlash SYBR Green qPCR Kit in the ABI Quant Studio 5 Sequence Detection System (Life Technologies, USA). The expression of the genes of interest was analyzed by the ΔΔCt method using ATP5G rRNA and GAPDH as internal control genes. Real-time primer information is provided in **Supplementary Table 9**.

### Cytosolic/nuclear fractionation

Cells were incubated with hypotonic buffer (25 mM Tris–HCl (pH 7.4), 1 mM MgCl_2_, 5 mM KCl, and RNase inhibitor) on ice for 5 min. An equal volume of hypotonic buffer containing 1% NP-40 was added, and the sample was left on ice for another 5 min. After centrifugation at 5,000*g* for 5 min, the supernatant was collected as the cytosolic fraction. The pellets were resuspended in nuclear resuspension buffer (20 mM HEPES (pH 7.9), 400 mM NaCl, 1 mM EDTA, 1 mM EGTA, 1 mM dithiothreitol, 1 mM phenylmethyl sulfonyl fluoride and RNase inhibitor) and incubated at 4 °C for 30 min. The nuclear fraction was collected after removing insoluble membrane debris by centrifugation at 12,000*g* for 10 min. RNA isolation was performed using the Trizol method.

### RNA stability assays

U87 cells were treated with 5 μg/ml actinomycin D at various times as indicated (0, 60, 120 minutes). RNA was extracted, cDNA was made, and qRT–PCR was carried out as described above. RNA half-life (t1/2) was calculated by linear regression analysis.

### ChIRP assay

The targets of LncRNA were identified using ChIRP-RNA sequencing, which was described earlier (Chu, Quinn and Chang, 2012). Antisense DNA probes for PITAR were designed using the Stellaris Probe Designer tool. Probes were labeled with Biotin-TEG at the 3′ ends. U87 cells were crosslinked with 1% glutaraldehyde for 10 min at 37 °C and then quenched with 0.125 M glycine buffer for 5 min. U87 cells were lysed in lysis buffer (50 mM Tris, pH 7.0, 10 mM EDTA, 1% SDS, DTT, PMSF, protease inhibitor, and RNase inhibitor) on ice for 30 min, and genomes were sonicated three times into fragments 300–500 bp in length. Chromatins were diluted twice the volume of hybridization buffer (750 mM NaCl, 1% SDS, 50 mM Tris, pH 7.0, 1 mM EDTA, 15% formamide, DTT, PMSF, protease inhibitor and RNase inhibitor). Biotin-TEG labeled probes (odd, even, and LacZ) were added, and mixtures were rotated at 37°C for four hours. Streptavidin-magnetic C1 beads were blocked with 500 ng/µl yeast total RNA and 1 mg/ml BSA for one hour at 25 °C and washed three times before use. We incubated biotin probes with U87 cell lysates and then used Streptavidin C1 magnetic beads for capture. Finally, beads were resolved for RNA by the RNA elution buffer. The eluted RNA was subjected to RNA sequencing. The raw reads were quality-analyzed to quantify the abundance of mRNA in the ChIRP assay. Upon satisfactory assessment, Kallisto was used with default parameters to quantitate and obtain TPM-normalized gene abundance. IGV was used to visualize the raw reads. The Biotin-TEG labeled Antisense probe sequences are provided in **Supplementary Table 9**.

### Immunoblotting and co-immunoprecipitation

The western blot analysis was described earlier (Fong et al., 2018). In brief, RIPA buffer with protease inhibitor cocktail was used to isolate protein from the GBM cell lines, and GSCs were quantified by Bradford’s reagent. These studies used the following antibodies: p53, p21, GAPDH, anti-Ub, and TRIM28. To measure the stability of the p53 protein, the cells were treated with Cycloheximide at a final concentration of 50µg/ml, and immunoblotting was performed from whole cell lysates.

The endogenous ubiquitination assay was described earlier (Fong *et al*., 2018b). Briefly, we transfect the HA-Ubiquitin in VC/PITAR OE stable U87 cells and performed co-immunoprecipitation with anti p53 antibody to detect the ubiquitination of the p53 protein; MG132 treated and untreated cells were lysed in IP lysis buffer (0.5% NP-40, 150 mM NaCl, 20 mM HEPES, pH 7.4, 2 mM EDTA, and 1.5 mM MgCl2 and 20mM Iodoacetamide (IAA)) supplemented with protease inhibitor cocktail for half an hour on ice. Cell lysates were incubated with protein G-Magnetic beads (Dynabeads Protein G beads) coated with the p53 antibodies overnight at 4°C, and then the IP products were washed three times with IP lysis buffer; after that, the proteins were eluted with SDS sample buffer, and the eluted protein were run on SDS–PAGE followed by immunoblotting and probed with anti-ubiquitin antibody.

### Luciferase reporter assay

Luciferase assays were performed using reporter lysis buffer (Catalog #E3971, Promega, U.S.A) and luciferase assay reagent according to the manufacturer’s instructions. Briefly, plasmids were transfected in the cells plated in 12 well plates. For determining the comparative luciferase activity of PITAR OE/ ΔPITAR OE plasmid was co-transfected with TRIM28 3’UTR luc plasmid/ ΔTRIM28 3’UTR luc plasmid. Cells were harvested after 48h of transfection, and lysates were made. The luciferase assays were performed using a luciferase assay substrate (Catalog #E151A, Promega, U.S.A), and luciferase readings were recorded using a luminometer (Berthold, Germany) using an equal quantity of protein measured by Bradford assay. ß-Gal assays were performed to normalize the transfection differences of the PG13 Luc assay, and RFP fluorescence intensity was measured to normalize TRIM28 3’UTR Luc activity.

### In vitro transcription

The plasmid DNA template is used for in vitro synthesis of biotinylated PITAR/ ΔPITAR sense, antisense, generated by PCR amplification. The forward primer contained the T7 RNA polymerase promoter sequence, allowing for subsequent in vitro transcription. PCR products were purified using the DNA Gel Extraction Kit (Thermo Fisher), and used as templates for synthesizing biotin-labeled probes. The reaction was carried out using RiboMAX™ Large Scale RNA Production System (T7) (Catalog #P1300, Promega, U.S.A) and biotin-11-UTP (Catalog #AM8451, Invitrogen, U.S.A) as per manufacturer’s instructions. Briefly, 2.5 µg of linear template DNA was added, and the reaction was incubated at 37^◦^C for 1.5 hours. After alcohol precipitation, the RNA was resuspended in 20 µl of nuclease-free water. Primers used for amplifying the region from PITAR OE construct with T7 promoter overhang are provided in **Supplementary Table 9**.

### Biotinylated RNA pulldown assay

The in vitro transcribed RNA was treated with DNase (Ambion) and purified with an RNeasy kit (Qiagen). Twenty pmol biotinylated RNA (PITAR-Antisense, PITAR-Sense, and ΔPITAR-Sense) was incubated with 1 mg whole lysate prepared from U87 cells in binding buffer [10 mM HEPES, pH 7.4, 100 mM KCl, 3 mM MgCl_2_, 5% glycerol, 1 mM DTT, Yeast transfer RNA (50ng/ml) and heparin (5 mg/ml) for 4 hr at 4°C. The biotinylated RNA-RNA complexes were pulled down by incubation with Dynabeads M-280 Streptavidin (Thermo Fisher Scientific) for 4 hr at 4^0^C. After brief centrifugation, bound RNA in the pulldown material was separated by elution followed by the TRI method. Afterward, RNA was converted to cDNA and quantified by using qRT-PCR. In these assays, a nonspecific transcript derived from the T7 promoter-luciferase construct and bead alone was added with lysate as a control for nonspecific binding.

### Antisense oligo blocking followed by ChIRP assay

To investigate the TRIM28 interaction site on the PITAR transcript, we performed a competition ChIRP assay. In brief, the fragmented glutaraldehyde fixed RNA was preincubated with biotin-less antisense oligo number three, located at the energetically favorable binding site of the TRIM28 transcript. Next, we incubated with a biotin-labeled odd antisense probe set and performed a ChIRP pulldown assay. The interacting RNA was eluted, and we performed cDNA preparation followed by qRT-PCR to detect the interacting TRIM28 transcript.

### Transfection of GSCs, neurosphere assay, and sphere diameter measurement

The transfection in GSCs was carried out in a single-cell suspension state for siRNA and plasmid DNA. 100 nM concentration of either Control non-targeting siRNA or gene-specific siRNA (Dharmacon, U.K), as indicated, were transfected using Dharmafect I (Dharmacon, U.K) according to the manufacturer’s instructions. Control vector or PITAR OE plasmids were transfected using Lipofectamine 2000 (Life Technologies, U.S.A) according to the manufacturer’s instructions. After 72 hours of transfection, cells were harvested and confirmed for gene manipulation by qRT-PCR. After 48 hours of transfection, the aggregates formed were dissociated into single cells, counted, and equal numbers of cells were plated at a density of 2 cells/µl in 24-well or 6-well plates. The number of spheres was counted after seven days of plating. Fresh medium was replenished every 2-3 days. Sphere diameter measurements were analyzed using ImageJ software. The number of spheres above 50µM diameter was counted and plotted for the total number of spheres.

### Limiting dilution assay

For each condition, GSCs (single cells) were plated 1, 10, 50, 100, and 200 cells in 10 wells each, respectively, of a 96-well plate, and sphere formation was assessed over the next 5– 7 days. The number of wells not forming spheres was counted and plotted against the number of cells per well. Extreme limiting dilution assay was done using the online ELDA software (https://bioinf.wehi.edu.au/software/elda/).

### Xenograft Orthotopic mouse model

U87MG-Luc cells bearing Control siNT/VC/shNT or siPITAR#1/PITAR OE/shTRIM28 (0.3 × 10^6^) were intracranially injected into the right corpus striatum by stereotactic injection 3 mm deep of 6-to 8-week-old immunocompromised CD1 nu/nu female mice (IISc, CAF; n = 10 mice per group). The mice were kept in a 12 hr light, and dark cycle, fed ad libitum with a regular diet, and the experiments were done in the light phase of the cycle. Intracranial tumors were monitored by bioluminescence imaging with the PerkinElmer IVIS Spectrum using mild gas anesthesia (using isoflurane) for the animals, and total photon flux (photons/s) was measured every 5-6 days intervals and plotted. Temozolomide (100mg/kg BW) was administered intraperitoneally every day for one week. After four weeks of tumor injection, mice were sacrificed. Studies on animals were conducted with approval from the Animal Research Ethics Committee of the IISc (IAEC approval No: CAF/Ethics/692/2019). The survival of mice in both groups (siNT and si PITAR#1, n=6) was followed up, and the survival curve was plotted using Graph pad Prism.

### RNAScope

RNAScope in-situ hybridization (ISH) was performed using RNAscope Multiplex Fluorescent Reagent Kit v2 (Cat. No. 323100) from Advanced Cell Diagnostics (ACD), which was described earlier (Mahale et al., 2022). RNA probes specific to PITAR and TRIM28 were designed by ACD’s made-to-order probes service. Manual RNAScope ISH protocol from Advanced Cell Diagnostics (ACD, 323100-USM) was followed to perform single or double RNA staining. RNAScope manual procedure involved sample fixation, sample pretreatment, subsequent probe hybridization, signal amplification, and ISH signal detection. RNA in-situ (RNAScope) and immunofluorescence imaging were performed on a Zeiss LSM 880 airy scan confocal microscope at the Centre for Cellular Imaging Facility. Most images were captured on 40X or 60X oil immersion objectives with the laser emitting 405, 488, 561, and 670 nM wavelengths, depending on the fluorophores used in the experiments. The 3D reconstruction was performed using Imaris microscopy image analysis software 9.8.2 version.

### Cell viability

The Vi-cell reagent Kit (Beckman Coulter) was used according to the manufacturer’s instructions. Briefly, cells were seeded at 1 × 10^5^ per well in 12-well plates overnight before treatment as desired. Cells are harvested and resuspended in 1ml 1X PBS. The viable cell count was done in the Vi-Cell cell viability Analyzer (Beckman Coulter).

### Apoptosis

Apoptotic cells were quantitated using the Annexin V Apoptosis Detection Kit (BD Biosciences). In brief, cells were washed twice with cold PBS and then resuspended in binding buffer at a concentration of 1 × 10^6^ cells per ml. 100 μl of the solution (1 × 10^5^ cells) was transferred to a 5-ml culture tube, and 5 μl of annexin V was added. After incubation at room temperature for 15 min in the dark, an additional 400 μl of binding buffer was added to each tube, and cells were analysed using a flow cytometer within 1 h (FACSVerse, BD Biosciences).

### Cell cycle analysis

Cells were fixed by 70% ethanol at -20^0^C overnight and spun down at 4,000 r.p.m. Cell pellets were resuspended in PBS containing 0.25% Triton X-100 and incubated on ice for 15 min. After discarding the supernatant, the cell pellet was resuspended in 0.5 ml PBS containing 10 μg/ml RNase A and 20 μg/ml propidium iodide stock solution and incubated at room temperature in the dark for 30 min. Cells were then subjected to analysis using a flow cytometer (FACSVerse).

### Colony formation assay

U87 and U343 cells were transfected with the PITAR siRNA and control siRNA. Twenty-four hours later, 1 × 10^3^ cells were cultured in a 6-well plate. Two weeks later, cells were fixed, stained with crystal violet, and photographed. The percentage and intensity of the area covered by crystal violet-stained cell colonies were quantified using the ImageJ software.

### Fluorescent immunohistochemistry

The FFPE tissue sections (5 μm thick) were de-waxed and rehydrated. Antigen retrieval was performed in a pressure cooker for 20 min in 10 mM Tris with 1 mM EDTA (pH 9). Nonspecific binding was blocked using blocking buffer (PBS (pH 7.4), 3% serum, 1% BSA, and 0.1% Tween) for 60 minutes at room temperature. Sections were then incubated with primary antibodies (Ki67, TRIM28, and p21) diluted in blocking buffer overnight at 4 °C. After washing twice with 0.1% PBS–Tween, slides were incubated with a secondary antibody conjugated with Alexa 488 (Thermo Fisher). After washing, sections were incubated with DAPI (Sigma-Aldrich) as a counterstaining. The sections were mounted using prolonged antifade glass mounting media (Thermo Fisher). Two investigators examined the slides. The percentage of positive cells was estimated from 0% to 100%. The intensity of staining (intensity score) was judged on an arbitrary scale of 0–4: no staining (0), weakly positive staining (1), moderately positive staining (2), strongly positive staining (3), and very strong positive staining (4). An immunoreactive score was derived by multiplying the percentage of positive cells with staining intensity divided by 10.

### H&E staining

Brain tissues were fixed with 4% paraformaldehyde, dehydration (gradient ethanol), and embedding in paraffin. Then, the Brain tissues were cut into 5-um slices using a microtome instrument. Afterward, the slices were dewaxed with xylene I and xylene II; dehydrated with 95%, 90%, 80%, and 70% ethanol; and addressed with distilled water. Finally, the slices were processed with Harris hematoxylin, 1% hydrochloric acid alcohol, 0.6% ammonia, and eosin. After dehydration (gradient ethanol) and immersion (xylene), the Brain tumor’s pathologic structure was observed with a microscope.

### Statistical analysis

Statistical analyses were performed using GraphPad Prism 6 (GraphPad Software, La Jolla, CA) or R software. Unpaired t-tests or one-way ANOVA, followed by t-tests for individual group comparisons (Tukey’s test), were used as described for each experiment. Data are presented as either means±s.e.m. or means±s.d. All the experiments were performed in biological triplicate unless otherwise specified. P < 0.05 was considered to be statistical significance.

## Data Availability

All high throughput data developed in this manuscript are provided in supplementary tables.

## Author contributions

SJ planned and carried out most of the experiments; MM, NJ, and AC helped in bioinformatics analysis; SM carried out RNA FISH; BG and LK carried out some aspects of genome analysis and RNA quantitation; KRP carried out part of animal studies and IHC; VA provided glioma tissue samples and corrected the manuscript; CK participated in RNA FISH and corrected the manuscript; KS prepare the manuscript, planned the study and lead the project.

## Acknowledgments

The results published here are in whole or part, based upon data generated by The Cancer Genome Atlas pilot project established by the NCI and NHGRI. Information about TCGA and the investigators and institutions that constitute the TCGA research network can be found at http://cancergenome.nih.gov/. We acknowledge the shRNA consortium (Dr. Subba Rao), IISc, India, for shRNA constructs. SJ acknowledges NPDF, DBT RA, and DST for fellowship. BG acknowledges DST Inspire, KRP acknowledges DBT for fellowship. KS acknowledges CEFIPRA, DBT, DST, ICMR, and CSIR (Govt. of India) for research grants. Infrastructure supported by DST FIST, DBT-IISc partnership program, and UGC is acknowledged. KS is awarded J. C. Bose Fellowship from DST.

