## Supplementary.pdf for "*PITAR*, a DNA damage-inducible Cancer/Testis long noncoding RNA, inactivates p53 by binding and stabilizing TRIM28 mRNA"

### Supplementary Figure legends

**Supplementary Figure 1: A.** The plot depicts one to one correlation between GSCs and their corresponding DGCs. **B.** The genomic track shows active histone marks (H3K27Ac) in GSCs over DGCs derived from the GSE54791 dataset using the IGV tools.

**Supplementary Figure 2: A.** Schematic of Odd probes, Even probes, and siRNAs location map with reference to the PITAR transcript. **B.** The cell proliferation was measured by MTT assay upon PITAR knockdown compared to control U343 cells. **C.** Viable cell count was measured using a viable cell counter following the Trypan blue method in U343 cells. **D.** Colony formation assay was performed upon PITAR knockdown condition compared to control U343 cells. **E.** Cell cycle analysis was carried out in PITAR-silenced U343 cells. **F.** The apoptotic cell death was measured by Annexin V/PI staining in PITAR-silenced U343 cells. Data are shown as mean ± SD (n=3). ****P*-value <0.001, ***P*-value <0.01, **P*-value <0.05.

**Supplementary Figure 3: A.** Genomic track for TRIM28 derived from ChIRP-RNA sequencing using Odd, Even, and LacZ antisense probes. **B.** A competition assay based on antisense oligonucleotide blocking was performed upon fragmented total RNA, preincubated with unlabelled PITAR antisense probe # 3, which is located close to the TRIM28 binding region on the exon 1 of PITAR, followed by pulldown with the PITAR biotinylated antisense (odd) probe set of PITAR. **C**. RNAScope images of co-localized signals were quantified using Imaris microscopy image analysis software 9.8.2 version.

**Supplementary Figure 4: clinical relevance of PITAR and TRIM28 association. A, B, C, & D.** Expression in log2 fold change of TRIM28 in Normal vs. GBM, derived from the CGGA, TCGA, Kamoun, and Gravendeel dataset. GlioVis was used to obtain the gene expression matrix, and an unpaired t-test was performed using GraphPad Prism v6. **E.** Expression in log2 fold change of TRIM28 in our lab patient cohorts is quantified using qRT-PCR. **F, G, H, I, & J.** The correlation analysis was performed between PITAR and TRIM28 in glioma samples from the CGGA, TCGA, Kamoun, and Gravendeel databases and our lab patient cohort. **K.** Expression in log2 fold change of TRIM28 was quantified in GSC and corresponding DGC from the GSE54791 database. **L.** The correlation analysis between PITAR and TRIM28 in GSC is derived from the GSE92459 database. **M.** TRIM28 protein expression in Normal vs. GBM, derived from the CPTAC database. **N.** TRIM28 protein expression was quantified using immunohistochemistry images from the Protein atlas. Data are shown as mean ± SD (n=3). ****P*-value <0.001, ***P*-value <0.01, **P*-value <0.05.

**Supplementary Figure 5: A.** The PG13-Luc activity was measured in PITAR-silenced/siNT U87 cells. **B.** Relative expression of PITAR, TRIM28, TP53, and CDKN1A was quantified by qRT-PCR in PITAR-silenced U87 cells compared to siNT. **C.** The protein expression of TRIM28, p53, and p21 was measured by immunoblotting in PITAR-silenced U87 cells compared to siNT. **D & E.** Cells were transfected with pcDNA3.1-PITAR (PITAR OE)/ empty vector control plasmid (pcDNA3.1) and harvested 48h post-transfection for qRT-PCR (PITAR, TRIM28, TP53, and CDKN1A) and immunoblotting with indicated antibodies (TRIM28, p53, and p21). GAPDH served as the control. **F.** The PG13-Luc activity was measured in U87/siPITAR and U87/siNT cells with exogenously overexpressed TRIM28 conditions. **G.** The PG13-Luc activity was measured in U87/shTRIM28 and U87/shNT cells with exogenously overexpressed PITAR conditions. Data are shown as mean ± SD (n=3). ****P*-value <0.001, ***P*-value <0.01, **P*-value <0.05.

**Supplementary Figure 6: A.** The relative expression of p53 was quantified in U87/shNT and U87/shp53 cells by qRT-PCR. **B.** The immunoblot represents the knockdown efficiency of two shp53 clones and the expression of TRIM28 in the p53 silenced condition. **B.** The colony formation assay was performed and quantified in siNT and siPITAR conditions in a p53 knockdown background. **D.** The relative expression of PITAR, TRIM28, CDKN1A, and MDM2 was quantified in siNT and siPITAR conditions in a p53 Knockdown background using qRT-PCR. Data are shown as mean ± SD (n=3). ****P*-value <0.001, ***P*-value <0.01, **P*-value <0.05.

**Supplementary Figure 7: Glioblastoma stem-like cell growth is induced by PITAR through p53 inactivation. A.** The bright field image represents RG5 sphere growth in the siNT/siPITAR#1 condition. **B & C.** RG5 sphere growth was quantified by the measurement of sphere count and sphere size using ImageJ (https://imagej.nih.gov/ij/download.html). **D.** Limiting dilution assay was performed in RG5 cells. The graph represents the percentage of wells without spheres as a function of the number of PITAR Knockdown cells compared to control cells (https://bioinf.wehi.edu.au/software/elda/). **E.** Relative expression of PITAR, TRIM28, TP53, and CDKN1A was quantified by qRT-PCR in PITAR-silenced RG5 cells compared to siNT. **F.** The bright field image represents RG5 sphere growth in PITAR OE compared to the vector control. **G & H.** RG5 sphere growth was quantified by measuring sphere count and size in the PITAR OE condition compared to the vector control using ImageJ software. **I.** Relative expression of PITAR, TRIM28, TP53, and CDKN1A was quantified by qRT-PCR in the PITAR OE condition of RG5 cells compared to the vector control. **J.** The bright field image represents MGG8 sphere growth on siNT/siPITAR condition. **K & L.** MGG8 sphere growth was quantified by the measurement of sphere count and sphere size using ImageJ tools. **M.** Limiting dilution assay was performed in MGG8 cells. The graph represents the percentage of wells without spheres as a function of the number of PITAR Knockdown cells compared to control cells. **N.** The silencing efficiency of PITAR in MGG8 cells was measured using qRT-PCR. Data are shown as mean ± SD (n=3). ****P*-value <0.001, ***P*-value <0.01, **P*-value <0.05.

**Supplementary Figure 8: A.** The PG13-Luc activity was measured in the presence and absence of Adriamycin in U87/PITAR OE and U87/VC cells. **B**. The relative expression of PITAR and TRIM28 was measured in the presence and absence of Adriamycin and CGK733 using qRT-PCR in U87 cells. **C.** The TRIM28 protein expression was measured by immunoblotting in the presence of Adriamycin and CGK733 in U87 cells. **D.** The TRIM28 3'UTR Luc activity was measured in the presence of Adriamycin and CGK733 in U87 cells. **E.** The U87/siNT and U87/siPITAR cells were *γ*-irradiated (7Gy), and cell extracts were prepared at different time points postirradiation as indicated. The immunoblotting for MDM2, p53, and p21 were performed to show their kinetics. **F, G & H.** The quantification for MDM2, p53, and p21 immunoblot was performed and plotted with nonlinear regression curve fit. Data are shown as mean ± SD (n=3). ****P*-value <0.001, ***P*-value <0.01, **P*-value <0.05.

**Supplementary Figure 9: A.** H&E staining was performed on formalin-fixed tumor-bearing (VC, PITAR OE, VC+TMZ, and PITAR OE+TMZ) mouse brain sections. **B.** Immunohistochemistry was performed to detect TRIM28, Ki67, and p21 protein expression in tumor sections derived from VC, PITAR OE, VC+TMZ, and PITAR OE+TMZ tumors. The green color represents specific protein staining, and the blue depicts the nucleus stained with DAPI. Magnification = ×20, scale bar = 50 μm. **C.** Immunohistochemistry was performed to detect TRIM28 and p21 protein expression in tumor sections derived from PITAR OE/shNT and PITAR OE/shTRIM28 tumors. The red color represents specific protein staining, and the blue depicts the nucleus stained with DAPI. Magnification = ×40, scale bar = 40 μm. **D.** Correlation analysis was performed in the p53 wild type and p53 mutant GBM patient cohort. **E.** Univariate Cox regression analysis was performed in the p53 wild type and mutant p53

GBM patient cohort. **F, G & H.** The Kaplan–Meier graph depicts the survival correlation of PITAR in the total GBM cohort, p53 wild type, and p53 mutant GBM patient cohort.
