## Supplementary Figures.pdf for "*PITAR*, a DNA damage-inducible Cancer/Testis long noncoding RNA, inactivates p53 by binding and stabilizing TRIM28 mRNA"

### Supplementary Figures (1 to 9)

A

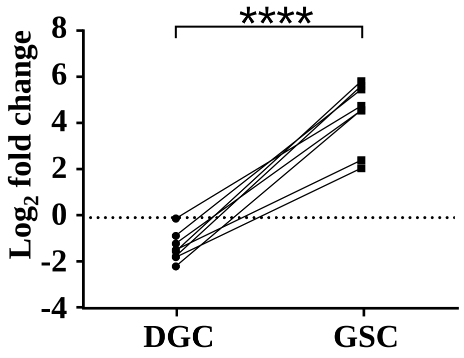
****
Log2 fold change
8
6
4
2
0
-2
-4
DGC
GSC

B

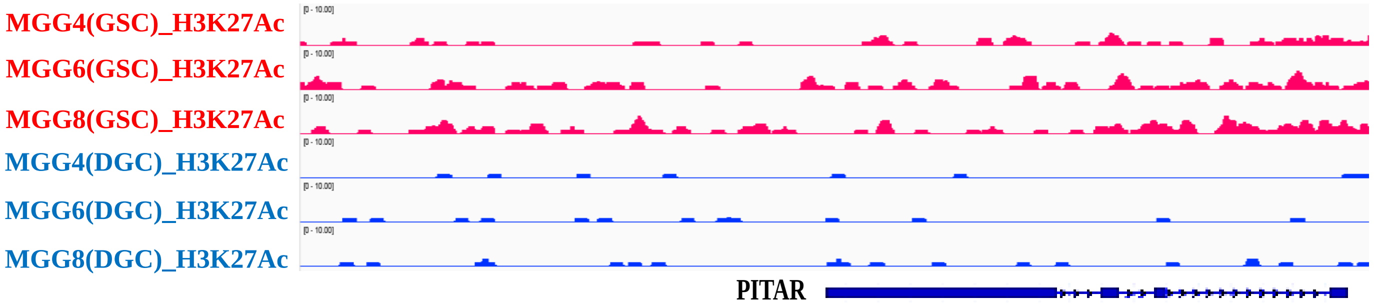
MGG4(GSC)_H3K27Ac
MGG6(GSC)_H3K27Ac
MGG8(GSC)_H3K27Ac
MGG4(DGC)_H3K27Ac
MGG6(DGC)_H3K27Ac
MGG8(DGC)_H3K27Ac
PITAR

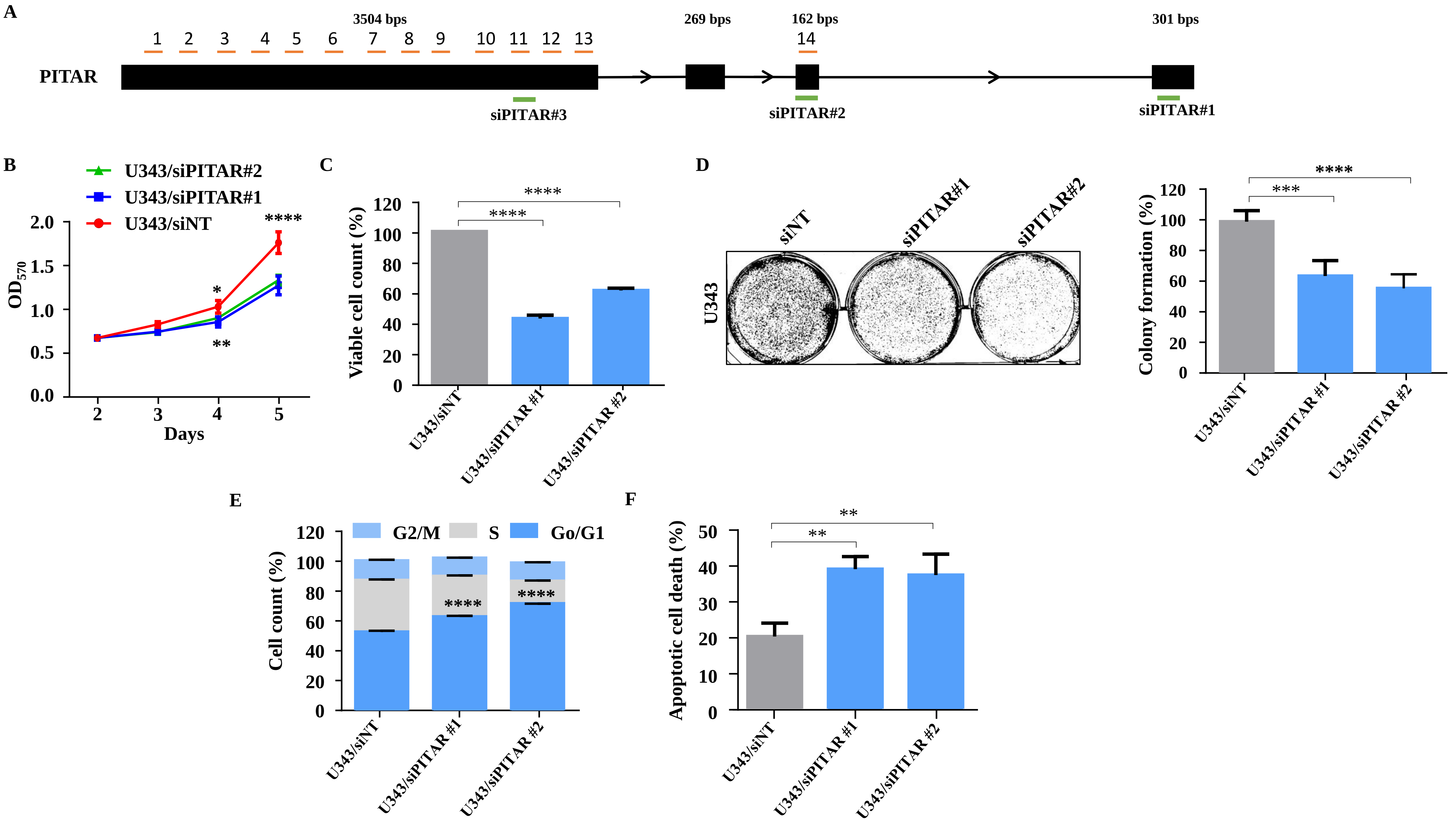
A
3504 bps
269 bps
162 bps
301 bps
1 2 3 4 5 6 7 8 9 10 11 12 13
14
PITAR
siPITAR#3
siPITAR#2
siPITAR#1
B
U343/siPITAR#2
U343/siPITAR#1
U343/siNT
$OD_{570}$
Days
C
Viable cell count (%)
U343/siNT
U343/siPITAR #1
U343/siPITAR #2
D
siNT
siPITAR#1
siPITAR#2
U343
Colony formation (%)
U343/siNT
U343/siPITAR #1
U343/siPITAR #2
E
G2/M
S
Go/G1
Cell count (%)
U343/siNT
U343/siPITAR #1
U343/siPITAR #2
F
Apoptotic cell death (%)
U343/siNT
U343/siPITAR #1
U343/siPITAR #2

A

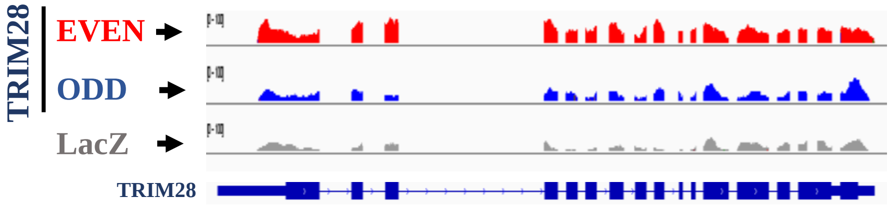
TRIM28
EVEN
ODD
LacZ
TRIM28

B

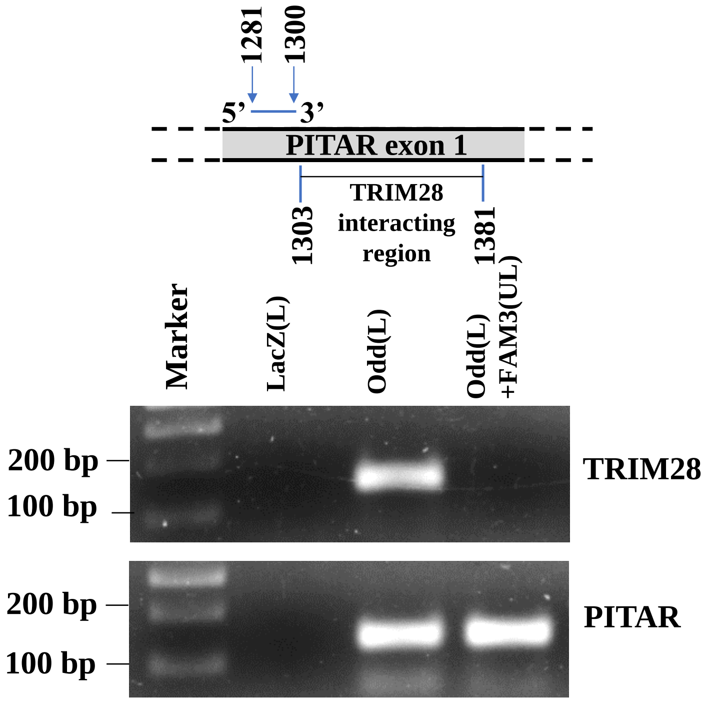
1281
1300
5'
3'
PITAR exon 1
TRIM28
interacting
region
1303
1381
Marker
LacZ(L)
Odd(L)
Odd(L)
+FAM3(UL)
200 bp
100 bp
TRIM28
200 bp
100 bp
PITAR

C

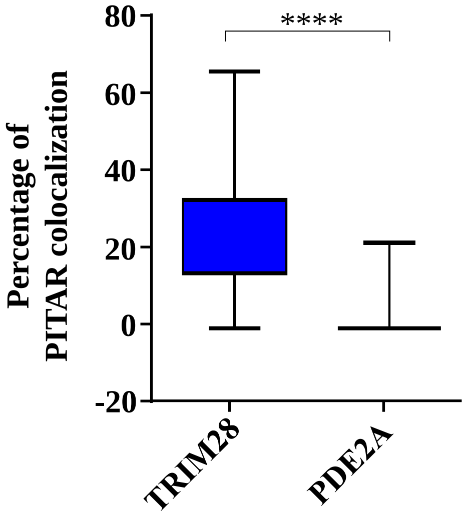
80
60
40
20
0
-20
****
Percentage of
PITAR colocalization
TRIM28
PDE2A

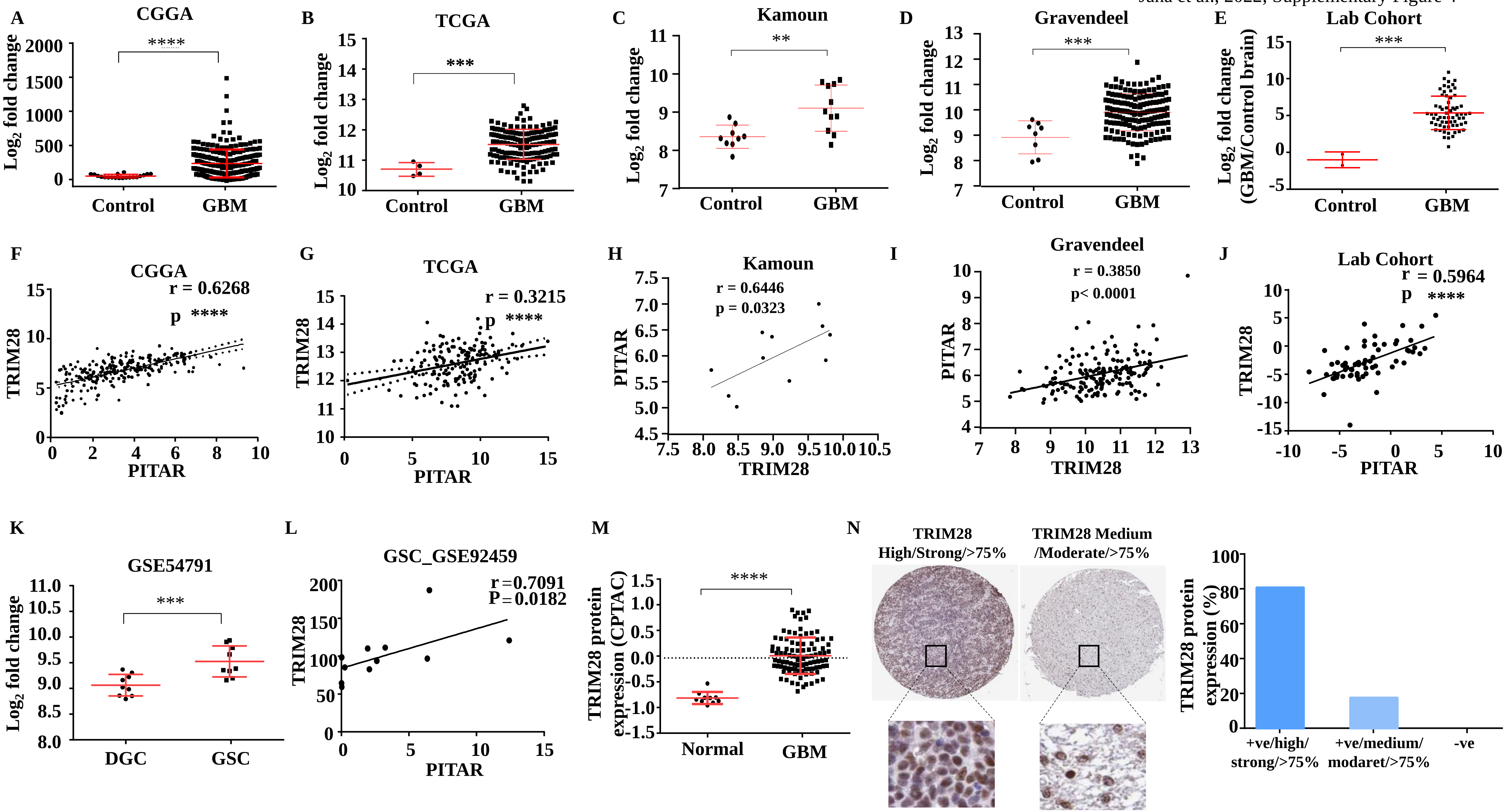
A
CGGA
****
Log2 fold change
Control
GBM
B
TCGA
***
Log2 fold change
Control
GBM
C
Kamoun
**
Log2 fold change
Control
GBM
D
Gravendeel
***
Log2 fold change
Control
GBM
E
Lab Cohort
***
Log2 fold change (GBM/Control brain)
Control
GBM
F
CGGA
r = 0.6268
p ****
TRIM28
PITAR
G
TCGA
r = 0.3215
p ****
TRIM28
PITAR
H
Kamoun
r = 0.6446
p = 0.0323
PITAR
TRIM28
I
Gravendeel
r = 0.3850
p< 0.0001
PITAR
TRIM28
J
Lab Cohort
r = 0.5964
p ****
TRIM28
PITAR
K
GSE54791
***
Log2 fold change
DGC
GSC
L
GSC_GSE92459
r = 0.7091
P = 0.0182
TRIM28
PITAR
M
****
TRIM28 protein expression (CPTAC)
Normal
GBM
N
TRIM28 High/Strong/>75%
TRIM28 Medium /Moderate/>75%
TRIM28 protein expression (%)
+ve/high/ strong/>75%
+ve/medium/ modaret/>75%
-ve

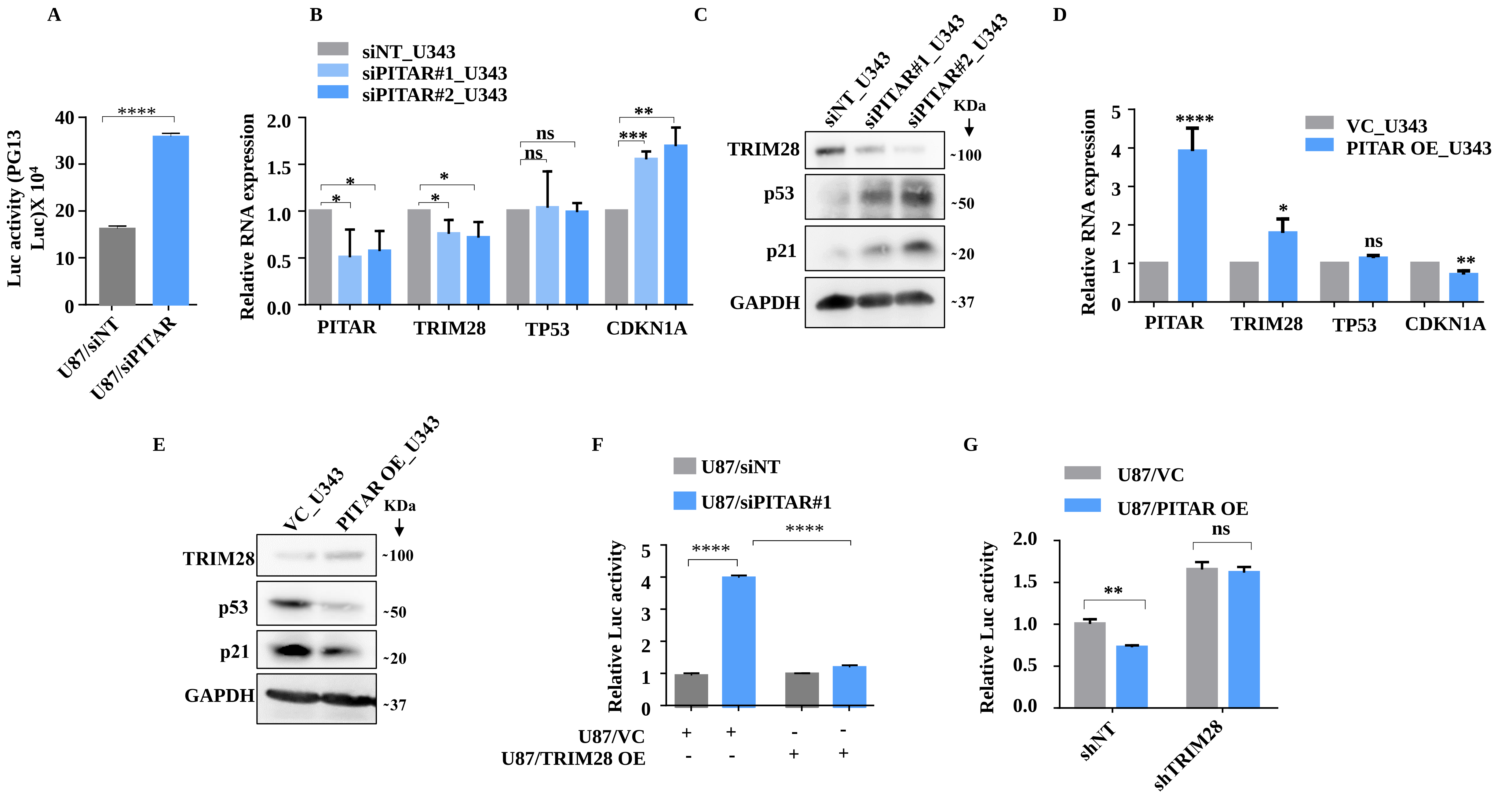
A
Luc activity (PG13 Luc)X 10^4
****
U87/siNT
U87/siPITAR
B
siNT_U343
siPITAR#1_U343
siPITAR#2_U343
Relative RNA expression
PITAR
TRIM28
TP53
CDKN1A
ns
C
siNT_U343
siPITAR#1_U343
siPITAR#2_U343
KDa
TRIM28
p53
p21
GAPDH
~100
~50
~20
~37
D
VC_U343
PITAR OE_U343
Relative RNA expression
E
VC_U343
PITAR OE_U343
F
U87/siNT
U87/siPITAR#1
Relative Luc activity
U87/VC
U87/TRIM28 OE
G
U87/VC
U87/PITAR OE
Relative Luc activity
shNT
shTRIM28

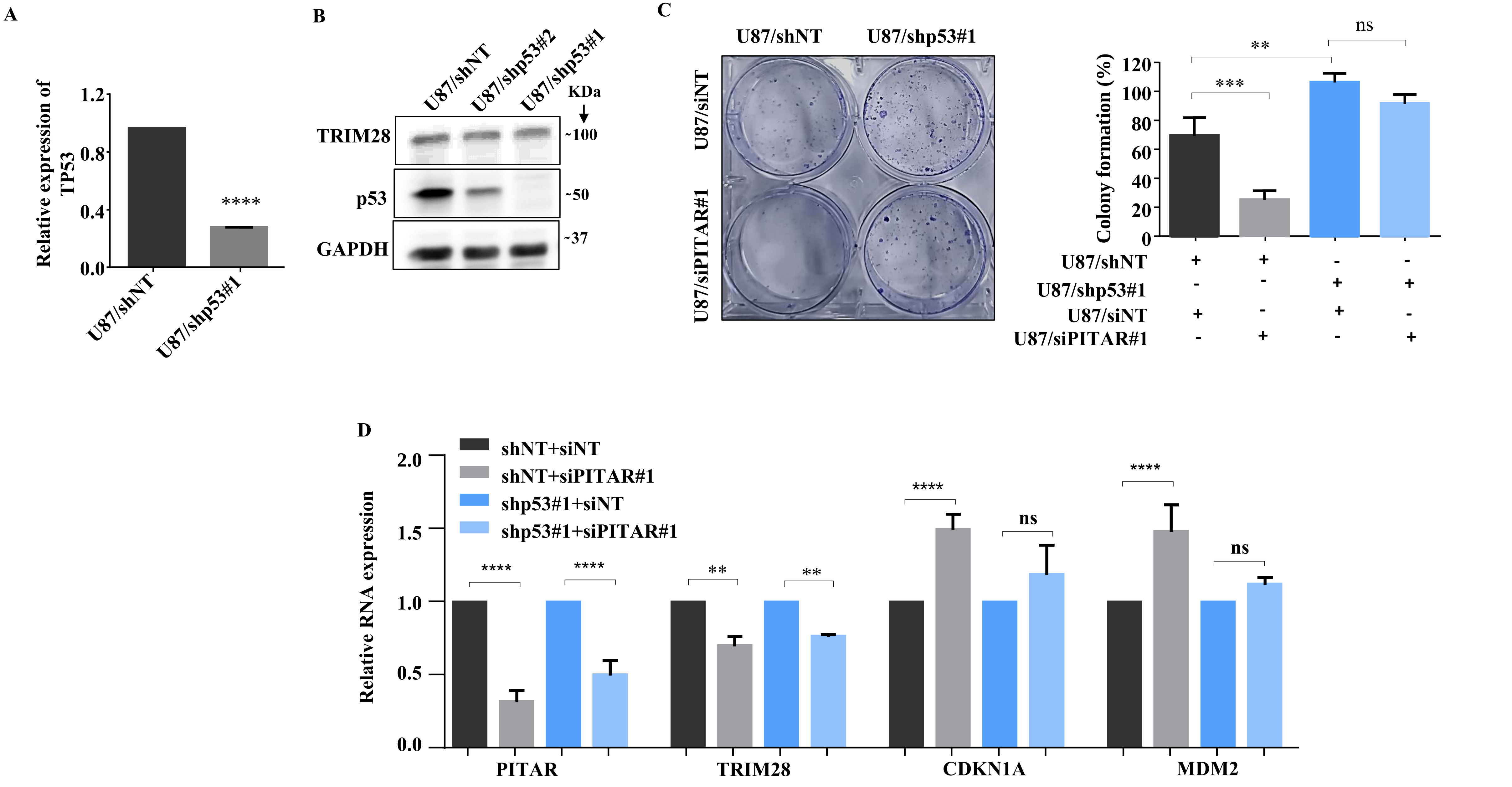
A
Relative expression of TP53
1.2
0.8
0.4
0.0
****
U87/shNT
U87/shp53#1
B
U87/shNT
U87/shp53#2
U87/shp53#1
KDa
TRIM28
~100
p53
~50
GAPDH
~37
C
U87/shNT
U87/shp53#1
U87/siNT
U87/siPITAR#1
Colony formation (%)
120
100
80
60
40
20
0
**
***
ns
U87/shNT + + - -
U87/shp53#1 - - + +
U87/siNT + - + -
U87/siPITAR#1 - + - +
D
shNT+siNT
shNT+siPITAR#1
shp53#1+siNT
shp53#1+siPITAR#1
Relative RNA expression
2.0
1.5
1.0
0.5
0.0
****
**
ns
PITAR
TRIM28
CDKN1A
MDM2

Jana et al., 2022; Supplementary Figure 7

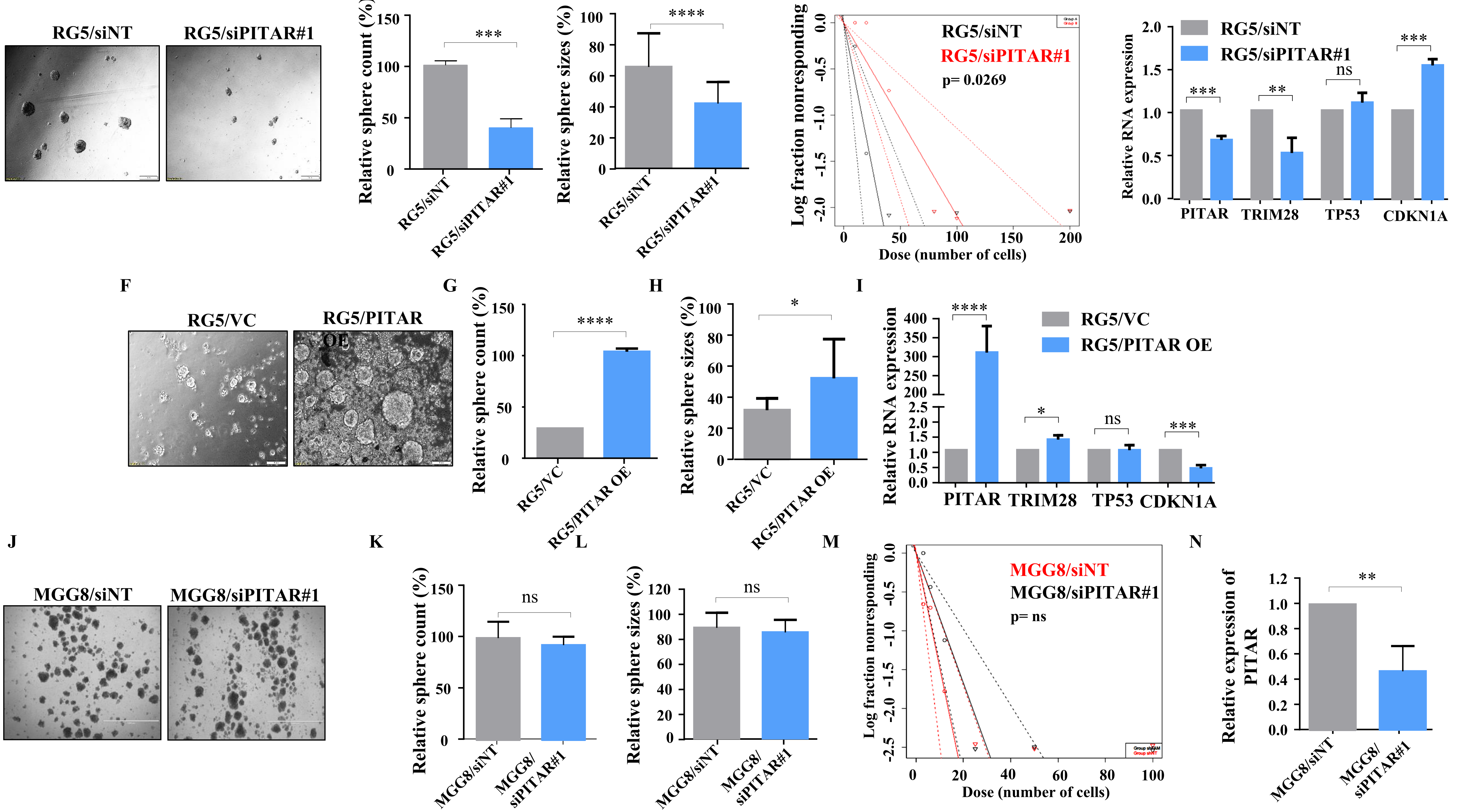

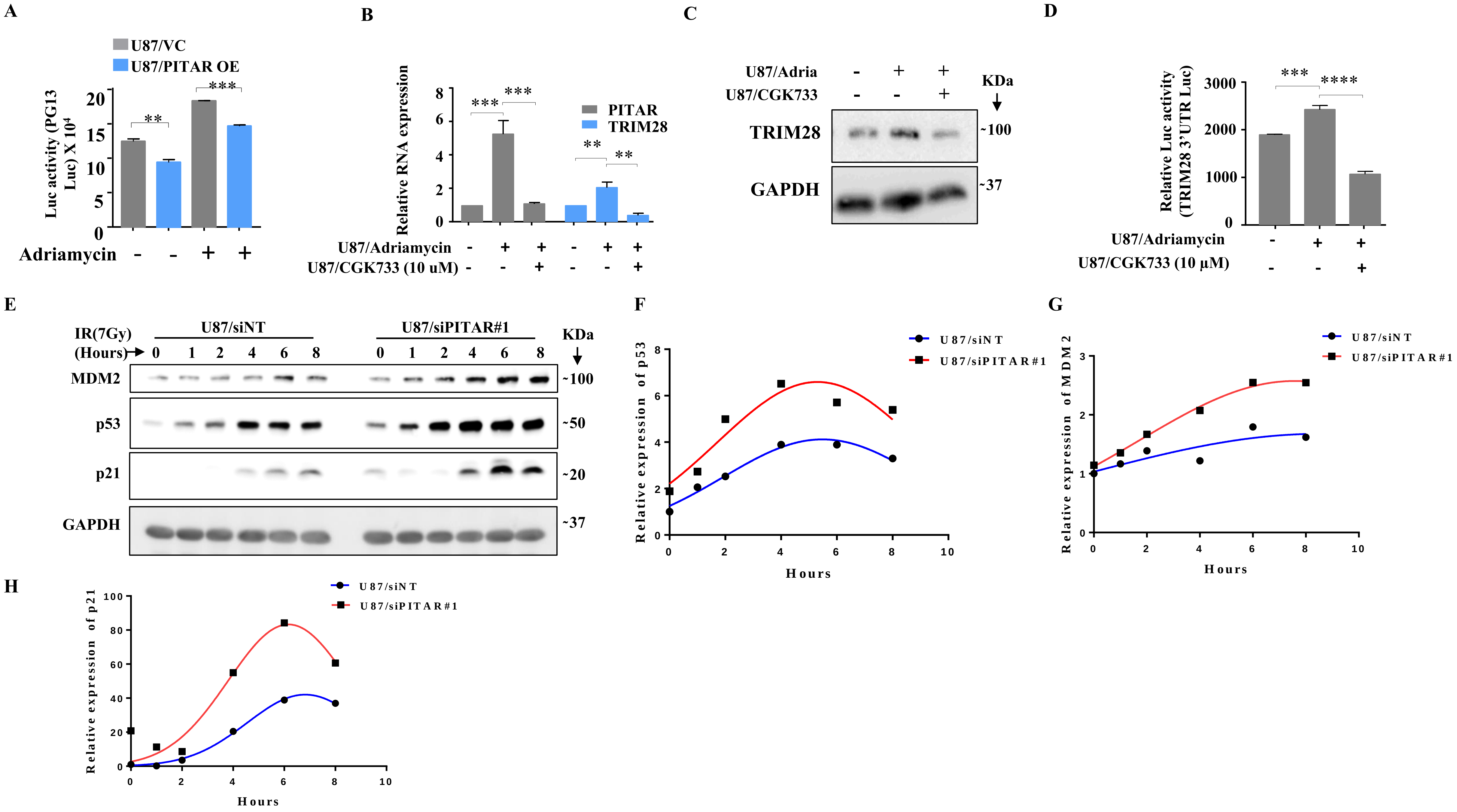
A
U87/VC
U87/PITAR OE
Luc activity (PG13 Luc) X 10^4
Adriamycin - - + +
B
Relative RNA expression
PITAR
TRIM28
U87/Adriamycin - + + - + +
U87/CGK733 (10 uM) - - + - - +
C
U87/Adria - + +
U87/CGK733 - - +
KDa
TRIM28 ~100
GAPDH ~37
D
Relative Luc activity (TRIM28 3'UTR Luc)
U87/Adriamycin - + +
U87/CGK733 (10 µM) - - +
E
IR(7Gy) (Hours)
U87/siNT
U87/siPITAR#1
0 1 2 4 6 8 0 1 2 4 6 8
KDa
MDM2 ~100
p53 ~50
p21 ~20
GAPDH ~37
F
Relative expression of p53
Hours
U87/siNT
U87/siPITAR#1
G
Relative expression of MDM2
Hours
U87/siNT
U87/siPITAR#1
H
Relative expression of p21
Hours
U87/siNT
U87/siPITAR#1

A

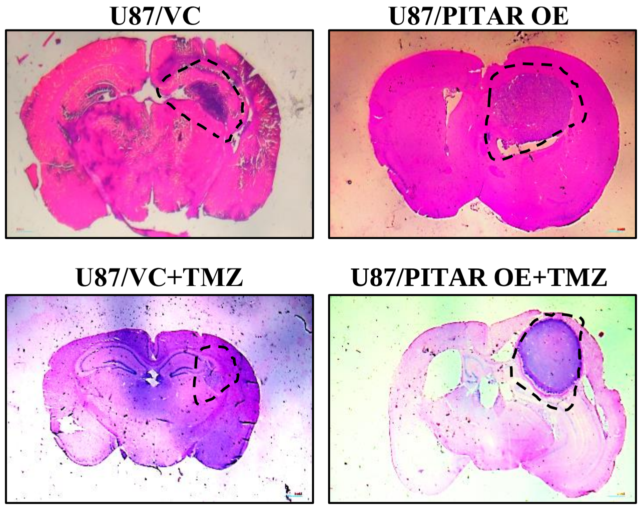


B

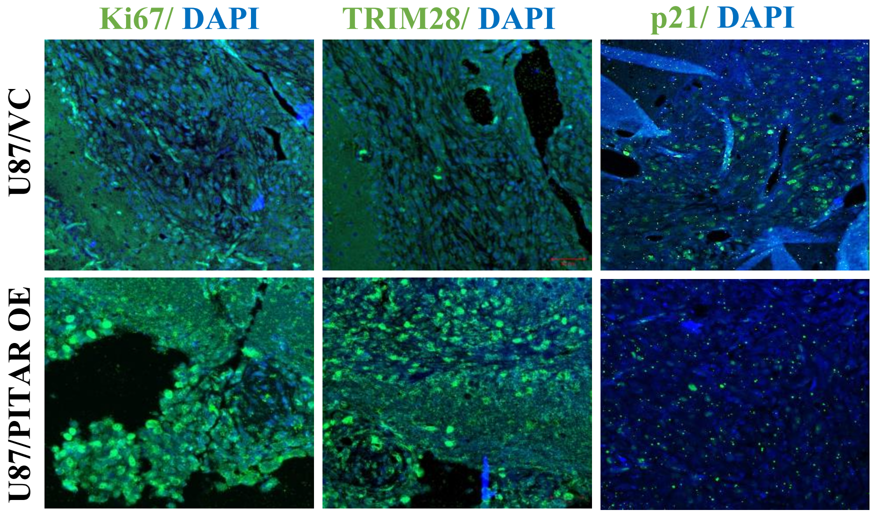


C

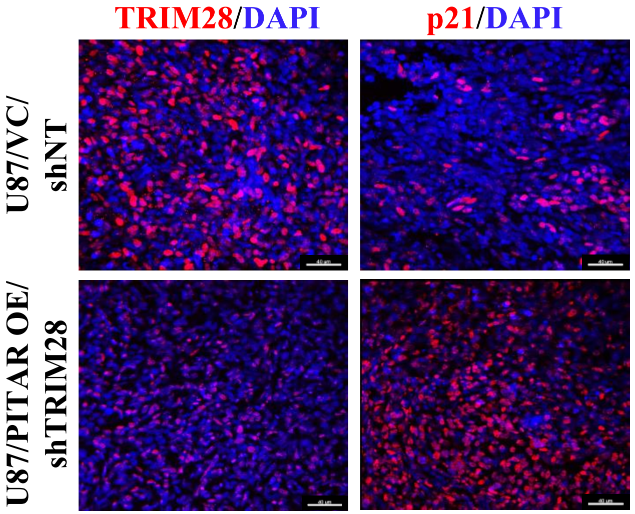


D

| Correlation analysis in 155 GBM patients | | | |
|---|---|---|---|
| Patient Cohort | Patient (n) | PITAR and TRIM28 | |
| | | r | p value |
| TCGA_GBM | 155 | 0.3215 | 0.0001 |
| TCGA_GBM_T53 (WT) | 101 | 0.3252 | 0.0009 |
| TCGA_GBM_T53 (MUT) | 54 | 0.2159 | 0.1169 |

E

| PITAR survival analysis in 155 GBM patients | | | | |
|---|---|---|---|---|
| Patient Cohort | Patient (n) | Univariate Analysis | | |
| | | Hazard Ratio | | |
| | | Estimate | 95% CI | p value |
| TCGA_GBM | 155 | 1.040 | 0.693-1.562 | 0.850 |
| TCGA_GBM_T53 (WT) | 101 | 1.655 | 1.018-2.688 | 0.042 |
| TCGA_GBM_T53 (MUT) | 54 | 0.633 | 0.297-1.348 | 0.236 |

F

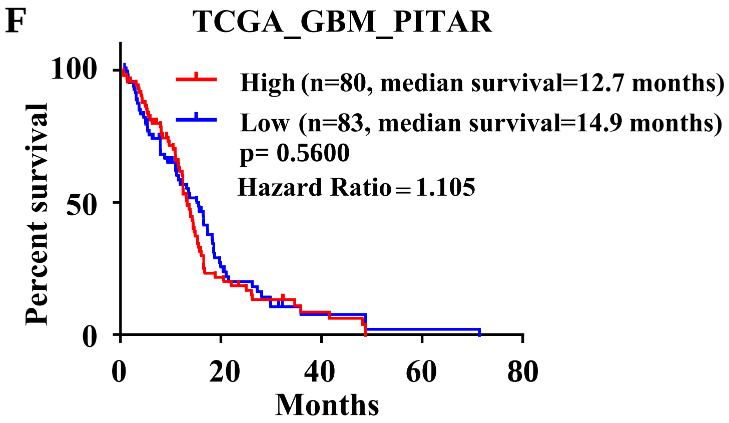


G

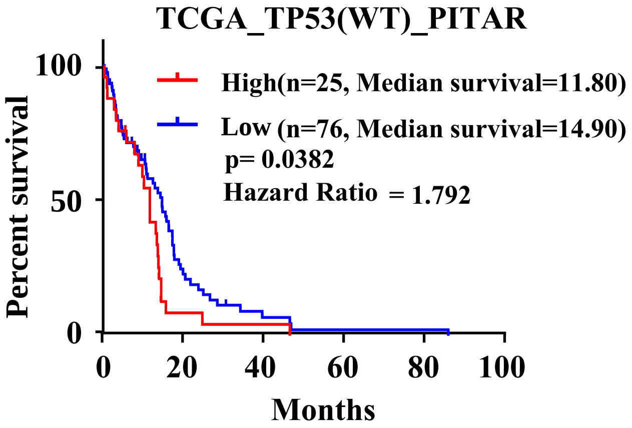


H

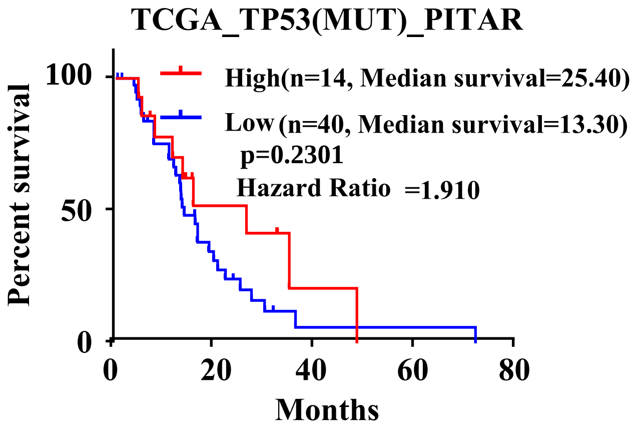
